# Responses of cortical neurons to intracortical microstimulation in awake primates

**DOI:** 10.1101/2022.03.30.486457

**Authors:** Richy Yun, Jonathan H. Mishler, Steve I. Perlmutter, Rajesh P. N. Rao, Eberhard E. Fetz

**Affiliations:** Department of Bioengineering, University of Washington; Department of Physiology and Biophysics, University of Washington; Department of Allen School for Computer Science and Engineering, University of Washington; Center for Neurotechnology, University of Washington; Washington National Primate Research Center, University of Washington

**Author notes:** Corresponding author: Richy Yun. Authors contributed equally (Richy Yun, Jonathan H. Mishler). The authors declare no competing financial interests.

## Abstract

Intracortical microstimulation (ICMS) is commonly used in many experimental and clinical paradigms; however, its effects on the activation of neurons are still not completely understood. To document the responses of cortical neurons in non-human primates to stimulation, we recorded single unit activity while delivering single-pulse stimulation via Utah arrays implanted in primary motor cortex of three macaque monkeys. Stimuli between 5-50 μA delivered to single channels reliably evoked spikes in neurons recorded throughout the array with delays of up to 12 milliseconds. ICMS pulses also induced a period of inhibition lasting up to 150 ms that typically followed the initial excitatory response. Higher current amplitudes led to a greater probability of evoking a spike and extended the duration of inhibition. The likelihood of evoking a spike in a neuron was dependent on the spontaneous firing rate as well as the delay between its most recent spike time and stimulus onset. Tonic repetitive stimulation between 2 and 20 Hz often modulated both the probability of evoking spikes and the duration of inhibition, although high frequency stimulation in particular was more likely to change both responses. On a trial-by-trial basis, whether a stimulus evoked a spike did not affect the subsequent inhibitory response; however, their changes over time were frequently positively or negatively correlated. Our results document the complex dynamics of cortical neural responses to electrical stimulation that need to be considered when utilizing ICMS for scientific and clinical applications.

**Significance statement:** Intracortical microstimulation (ICMS) is commonly used to probe the cortex, and previous studies have characterized the responses of single neurons to ICMS. However, these studies typically explored the averaged effects of ICMS throughout each experimental session, rather than by a trial-by-trial basis for each stimulation pulse. By shifting the approach, we explored the dependence of neural responses to ICMS on the spontaneous neural activity as well as the dynamics of responses over time due to repetitive stimulation. Our results highlight how the responses of neurons to ICMS are likely the result of interactions between local excitatory and inhibitory cortical circuits. These results will help inform the design of ICMS for both basic research and clinically relevant stimulation protocols.

## Introduction

Intracortical microstimulation (ICMS) is widely used for interfacing with the brain in both basic and clinical research, from inducing plasticity to employing sensory neuroprostheses in various animal models (Flesher et al., 2016; Hartmann et al., 2016; Jackson & Fetz, 2011; Lebedev & Nicolelis, 2017). The applicability of ICMS arises from the fact that it has the highest spatial and temporal specificity of all clinically applicable cortical stimulation techniques (Sejnowski et al., 2014). However, the circuit mechanisms that drive the responses of neurons following ICMS, and the ways in which other factors such as timing and stimulation frequency affect the stimulus responses are not fully understood.

ICMS was originally thought to activate neural elements around the electrode tip. Regions closer to the tip would have higher activation in a sphere with an isotropic gradient, and the volume would grow with increasing current amplitude (Ranck, 1975; Stoney et al., 1968; Tehovnik et al., 2006). However, it is now accepted that ICMS predominantly excites axons near the electrode tip that transsynaptically excite neurons up to several millimeters away, favoring pathways with similar functional properties (Gustafsson & Jankowska, 1976; Hao et al., 2016; Histed et al., 2009; Lesser et al., 2008; Logothetis et al., 2010; McIntyre & Grill, 2000). Additionally, the effects of ICMS are not limited to excitation, and includes a long-lasting inhibitory response that is commonly attributed to GABAergic synapses (Berman et al., 1991; Butovas & Schwarz, 2003; Hao et al., 2016).

Single neuron responses to ICMS are dynamic and can be modulated with repeated stimulation. The changes, in part, also depend on stimulus frequency. In particular, the excitation of neurons generally decreases over time and becomes more localized with higher frequencies (Dadarlat et al., 2019; Michelson et al., 2019). However, the reported frequency ranges and timescales are variable, and the driving processes remain unclear. The changes over time are often attributed to short-term synaptic plasticity or intrinsic plasticity of neurons which both depend on the frequency and pattern of stimulation (Abbott & Regehr, 2004; Citri & Malenka, 2008; Zucker & Regehr, 2002).

Altogether, these studies demonstrate that the effects of ICMS are not restricted to regions proximal to the electrode tip, and that responses consist of interplay between excitation and inhibition (Borchers et al., 2012; Griffin et al., 2011; Logothetis et al., 2010). Despite our increasing understanding of how ICMS activates cortical circuits, several significant questions remain. How does the background neuronal activity, including firing rate and previous spike time impact the stimulus response? How do the responses change over time as a function of both the frequency of stimulation and proximity to stimulation site? Is the inhibitory response coupled to the excitatory response, or are they independently activated?

We addressed the questions above by delivering ICMS and examining responses of single neurons in primary motor cortex (M1) of three macaque monkeys with chronically implanted Utah arrays. Single-pulse ICMS was delivered to one channel for up to 20 minutes while the spikes of single neurons were simultaneously recorded from all other electrode channels. We tracked the probability of evoking spikes as well as the duration of the evoked inhibition and varied both the sites and frequency of the stimulation between sessions. Our results expand upon previous findings by characterizing the dependencies of the neuronal responses to background neuronal activity, distance, and stimulation frequency, and exploring the interactions between the excitatory and inhibitory responses.

## Methods and Materials

### Experimental Design

#### Implants and surgery

Three pigtail macaque monkeys (*Macaca nemestrina*) were unilaterally (right hemisphere) or bilaterally implanted with 96-channel Utah microelectrode arrays (Blackrock Microsystems; 10×10, 400μm inter-electrode distance, 1.5 mm depth, Iridium oxide) in the hand region of M1. Sterile surgeries were performed under isoflurane anesthesia and aseptic conditions with continuous monitoring of all vitals. Animals received postoperative courses of analgesics and antibiotics following each surgery. All procedures conformed to the National Institutes of Health *Guide for the Care and Use of Laboratory Animals* and were approved by the University of Washington Institutional Animal Care and Use Committee.

Implantation of the arrays was guided via stereotaxic coordinates. A 1.5 cm wide square craniotomy centered at 4 mm lateral of Bregma was performed to expose the dura. Three sides of the exposed dura were cut to expose the cortex, after which a Utah microelectrode array was implanted. Two reference wires were inserted under the dura and two were inserted between the dura and the skull. The dura was then sutured around the implant, and the bone flap from the craniotomy was reattached to the skull with a titanium strap and titanium skull screws. A second, smaller titanium strap was screwed onto the skull to secure the wire bundle that connected the array to a connector “pedestal” that was also secured to the skull with eight skull screws. The skin incision was then sutured around the pedestal base.

To facilitate the chronic recording of neuronal activity, the monkeys were also implanted with halos made with 3/8” aluminum bars in an egg-shaped oval that was 17 cm long and 15.3 cm wide. Four titanium straps were affixed to the skull via titanium bone screws. Two of the straps were implanted bilaterally over the occipital ridge, and two were bilaterally implanted temporally. After the plates integrated with the skull for 6 weeks, the halo was secured to the skull with four pins, each of which were seated in one of the four plates.

#### Electrophysiology

Stimulation and recording of single unit activity were conducted with one of three systems: 1) Neurochip3 (custom bidirectional brain-computer interface developed in our laboratory (Shupe et al., 2021), 32 channels, 20 kHz sampling rate), 2) Neural Interface Processor (Ripple Neuro, 96 channels, 30 kHz sampling rate), or 3) RZ2 BioAmp Processor, PZ5 NeuroDigitizer Amplifier, and IZ2 Electrical Stimulator (Tucker-Davis Technologies, 96 channels, 25 kHz sampling rate).

#### Experimental Setup

The monkeys were trained to calmly sit in a primate chair while periodically receiving an apple smoothie reward without performing a task (Figure 1). Each session included a prestimulus epoch lasting between 5 and 10 minutes and a stimulus epoch lasting between 5 and 20 minutes. During the stimulus epoch we delivered tonic or Poisson distributed single pulse stimulation (cathodal, biphasic, 200µs phase width) to a single channel at rates between 1-20 Hz. For testing the effects of current amplitude, a range of 2-50 µA was used. Current amplitude was fixed at 15 µA for all other experiments and analyses. The stimulation frequency was fixed during the stimulus epoch for each session.

**Figure 1.**
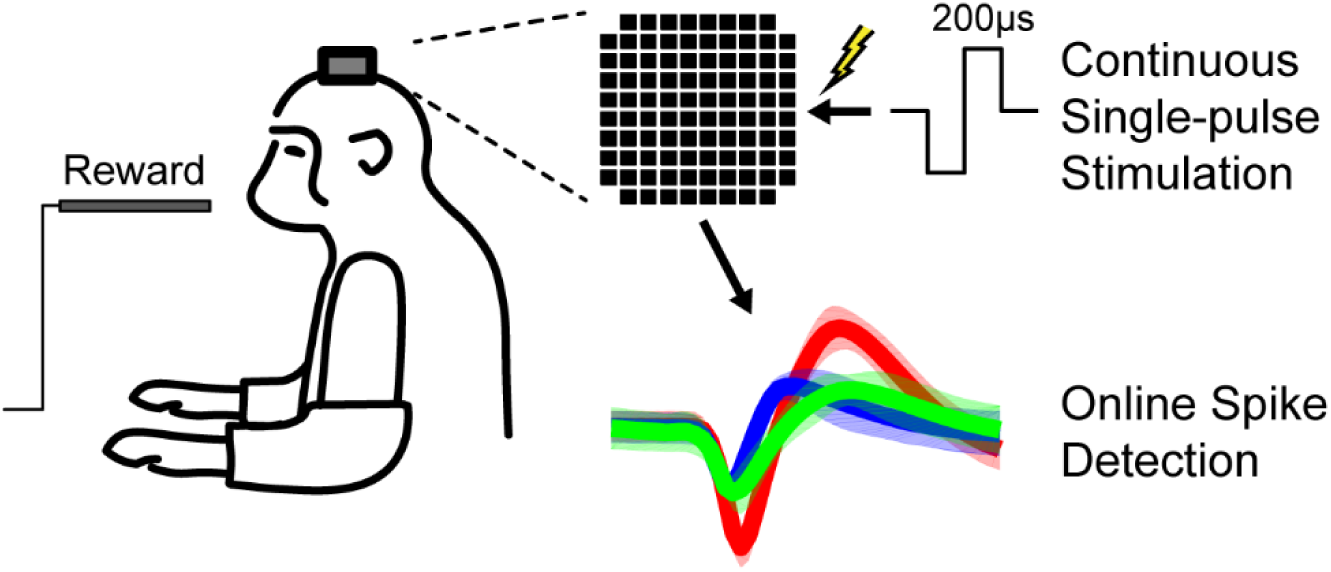
Experimental setup and timeline. Macaques calmly sat in a chair receiving apple smoothie reward through the experiment. Cathodic, 200us phase width, single-pulse ICMS was delivered to one channel of the Utah array in primary motor cortex while unit responses were recorded across the array. Each session consisted of a prestimulus and stimulus epoch.

### Data analysis

#### Evoked spike acquisition

Spikes were sorted using two time-amplitude windows, initially online and subsequently confirmed offline. Stimulus artifacts lasted around 1.1-1.6 ms. Spikes were frequently detected immediately following the artifact (Figure 2A). The timing of evoked spikes was found by calculating the peristimulus time histogram (PSTH, 0.5 ms bin widths) of spikes in the window from -20 to 20 ms from the time of stimulation (Figure 2B). To isolate the evoked spikes from the spontaneous activity, we defined upper and lower thresholds in the PSTH as the histogram mean plus or minus 2 times the standard deviation from -20 to -2 ms. We then found the largest peak in the PSTH from 1 to 15 ms after stimulation that was larger than the upper threshold and tracked adjacent bins in both directions until we reached the lower threshold on both sides. All spikes occurring within this window were denoted as stimulus-evoked spikes (Figure 2C). If no peak was greater than the threshold the spike was not considered to have been evoked by stimulation. The probability of evoked spikes was calculated as the number of evoked spikes divided by the number of total stimuli. For any analysis over time, the evoked spike probability was calculated for stimuli within overlapping 30 second bins with 1 second steps.

**Figure 2.**
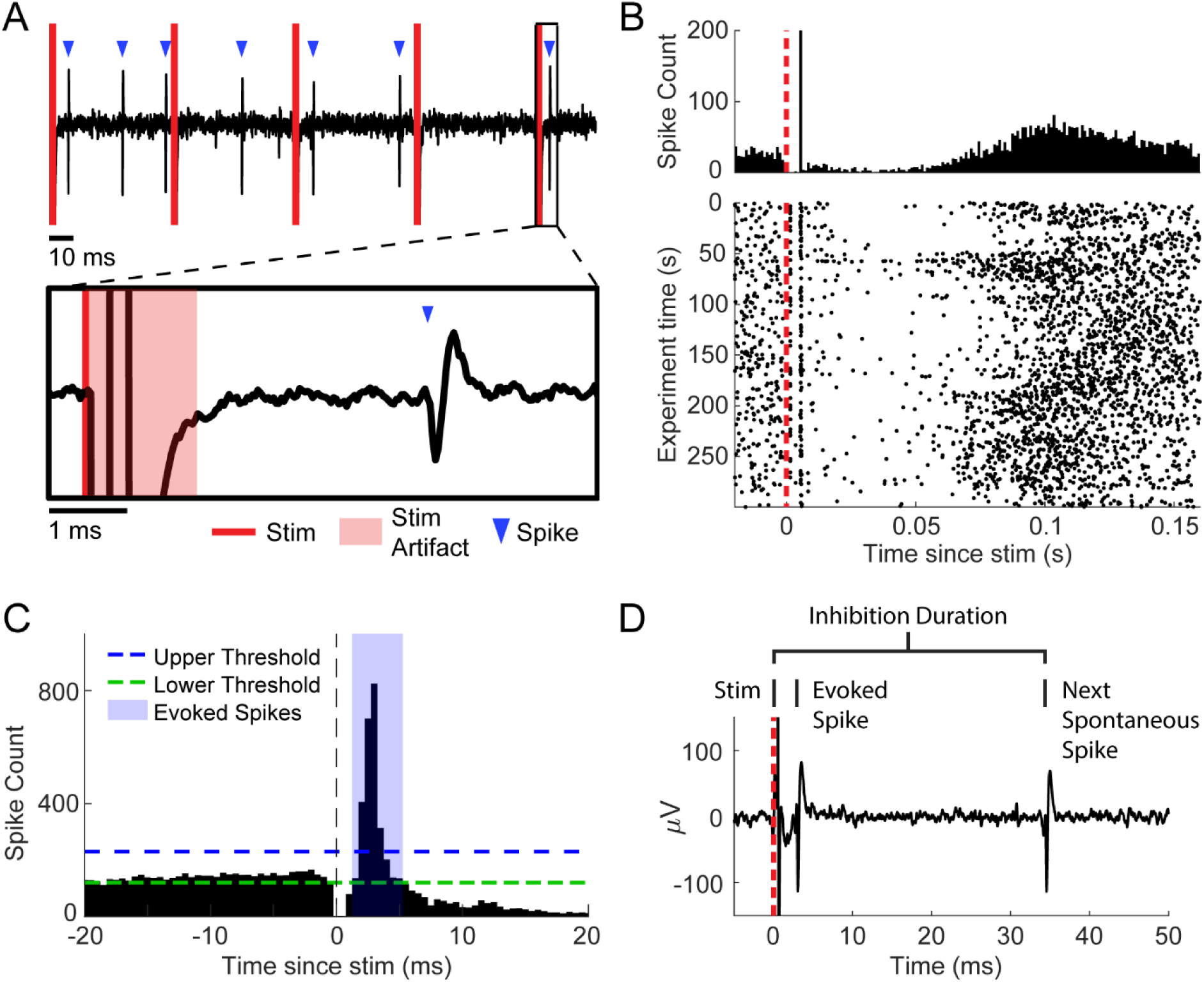
Detection of evoked spikes and inhibition. **A.** Example of filtered data trace. The inset shows a stimulus followed by an evoked spike after 3.5 ms. **B.** Example PSTH (top) and corresponding raster plot (bottom). **C.** Defining evoked spikes. A PSTH with 0.5 ms bins was generated. Peaks after the time of stimulation greater than the upper threshold (mean+2 standard deviations of -20 to -2ms in the PSTH) down to the lower threshold (mean-2 standard deviations) were called evoked spikes. **D.** Defining inhibition. Rather than using the PSTH, the inhibition duration was calculated for each stimulus by taking the time from stimulus onset to the next spontaneous spike.

We tracked a total of 148 distinct neurons across the array. Because of changes in signal fidelity due to electrode drift, we were not able to record all neurons across all sessions. As a result, the data presented in this study considers a “unit” to be one neuron recorded in one session. Across all sessions we recorded 585 units, 17 of which were exclusively used to analyze the effects of stimulus timing on the probability of evoking a spike since the analysis required high firing and/or stimulation rates. All units were used for all other analyses.

#### Inhibitory response acquisition

The inhibitory response was measured using the PSTH in previous studies (Butovas & Schwarz, 2003; Hao et al., 2016). While evoked spike timing can easily be determined with the PSTH due to their high probability and narrow time window, the inhibitory response depends on a broad window with sparse activity, particularly for units with low firing rate. As a result, a large number of stimuli is required to reliably detect inhibition in the PSTH. Therefore, rather than using the PSTH, we measured the duration of inhibition by removing all evoked spikes and calculating the time between the onset of stimulation and the next spontaneous spike (Figure 2D). Inhibition was deemed to be stronger when the delay from stimulation onset to the next spontaneous spike was longer, giving us a measure directly comparable to the PSTH but with much higher temporal resolution. We discarded any stimuli for which the subsequent stimuli occurred before the next spontaneous spike. For any analysis over time, we used the median inhibition duration of stimuli within overlapping 30 second bins with 1 second steps.

#### Evoked spike probability dependencies

To calculate the spontaneous firing rate for each unit over time, we disregarded the times from each stimulus onset to the next spontaneous spike. This effectively removed the stimulus response from the firing rate calculation, providing us with an independent measure of spontaneous activity.

The autocorrelation histograms for the 17 units in which we characterized the evoked spike probability as a function of the time delay between their previous spike times and stimulation onsets were calculated from their respective prestimulus epochs. The histograms were binned between 0 to 50 ms in non-overlapping 1 ms bins.

The dependencies of evoked spike probabilities on the timing of stimulation were calculated by first measuring the delays between each stimulus and the preceding spike. We separated this into two groups corresponding to whether the preceding spike was spontaneous or evoked. Stimuli that did not have a spike following the previous stimulus were discarded from the analysis. To account for the transmission delay from the stimulated site to the evoked spike, we added the average evoked spike latency for each unit to the delays. We then calculated the probability of evoking a spike for stimuli with delays from 0 to 50 ms with 1 ms time bins, then applied a moving average with a 5 ms window. Since the average spike latencies were added to each delay, time bins that were less than the average evoked spike latency were not included in correlation calculations.

#### Changes over time

Pairwise correlations and their statistical significance between firing rate, evoked spike probability, and inhibition duration over time were calculated using the Pearson correlation coefficient *r*:

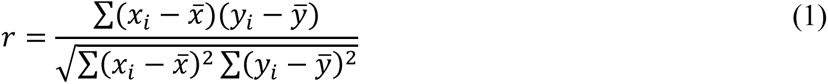

where 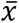 is the mean of the x-variable, and 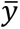 is the mean of the y-variable. Firing rate was calculated by removing all times between each stimulus onset to the first spontaneous spike (the inhibitory period) to remove any confounding affects between firing rate and the inhibitory response.

Linear and exponential fits were performed on binned evoked spike probabilities and inhibition durations to determine changes over time due to repetitive stimulation:

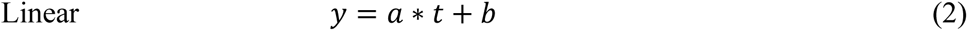

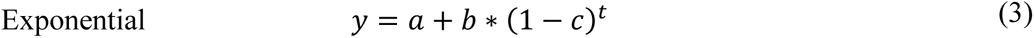

where *y* is either the evoked spike probability or inhibition duration, *t* is time and *a*, *b*, and *c* are the fitted variables. Changes were denoted to occur if the analysis of variance (ANOVA) F-statistic resulted in p<0.05. The sign of the linear fit slope (*a* in Equation 2) or the sign of the exponent base (*b* in **Error! Reference source not found.**3) of the exponential fit determined whether the changes were classified as increasing or decreasing. Neuron-stimulation site pairs with less than 3% average probability of evoking spikes were disregarded for analyses over time due to their inconsistency. The changes over time in spikes and inhibition were designated to be correlated if their correlation had a p-value less than 0.05.

#### Statistical analysis

Two-sided Wilcoxon signed-rank test (*signrank*, MATLAB) was used to compare between groups due to the nonparametric nature of the data. Two-sided Wilcoxon rank-sum (*ranksum*, MATLAB) tests were used for paired data. Fisher’s exact test (*fishertest*, MATLAB) or two-way analysis of variance (ANOVA) (*anova2*, MATLAB) was used to compare categorical data. The Pearson correlation was used for all correlation tests. The Kolmogorov-Smirnov test was used to compare distributions. P-values for significance and tests used are reported in individual analyses.

## Results

### Evoked spikes and inhibitory response

We found that ICMS elicited a brief excitatory response followed by a longer inhibition. Electrodes on the Utah array typically showed evoked spikes occurring 1.5 to 10 ms after single-pulse stimulation. The inhibition typically followed the excitatory response and was observed as the suppression of firing in the PSTH for 5 - 100 ms, although in some instances it lasted up to 200 ms. We also observed that stimulus-evoked inhibition could occur in the absence of the excitatory response.

To ensure that stimulation arriving during inhibition was not affecting the stimulus response, we delivered trains of two or three pulses with each subsequent pulse timed to occur during the inhibitory response of the previous stimulus. Single, double, and triple pulse stimuli were delivered to a single channel, randomly interleaved at 2 Hz. Our results across 5 different sessions show that stimuli delivered during the inhibitory response were able to reliably evoke spikes comparable to when stimuli was delivered at other times, as previously reported (Butovas & Schwarz, 2003) (Figure 3, left). Furthermore, each stimulus pulse “reset” the inhibitory response such that the duration of inhibition was the same following each pulse train (Figure 3, right).

**Figure 3.**
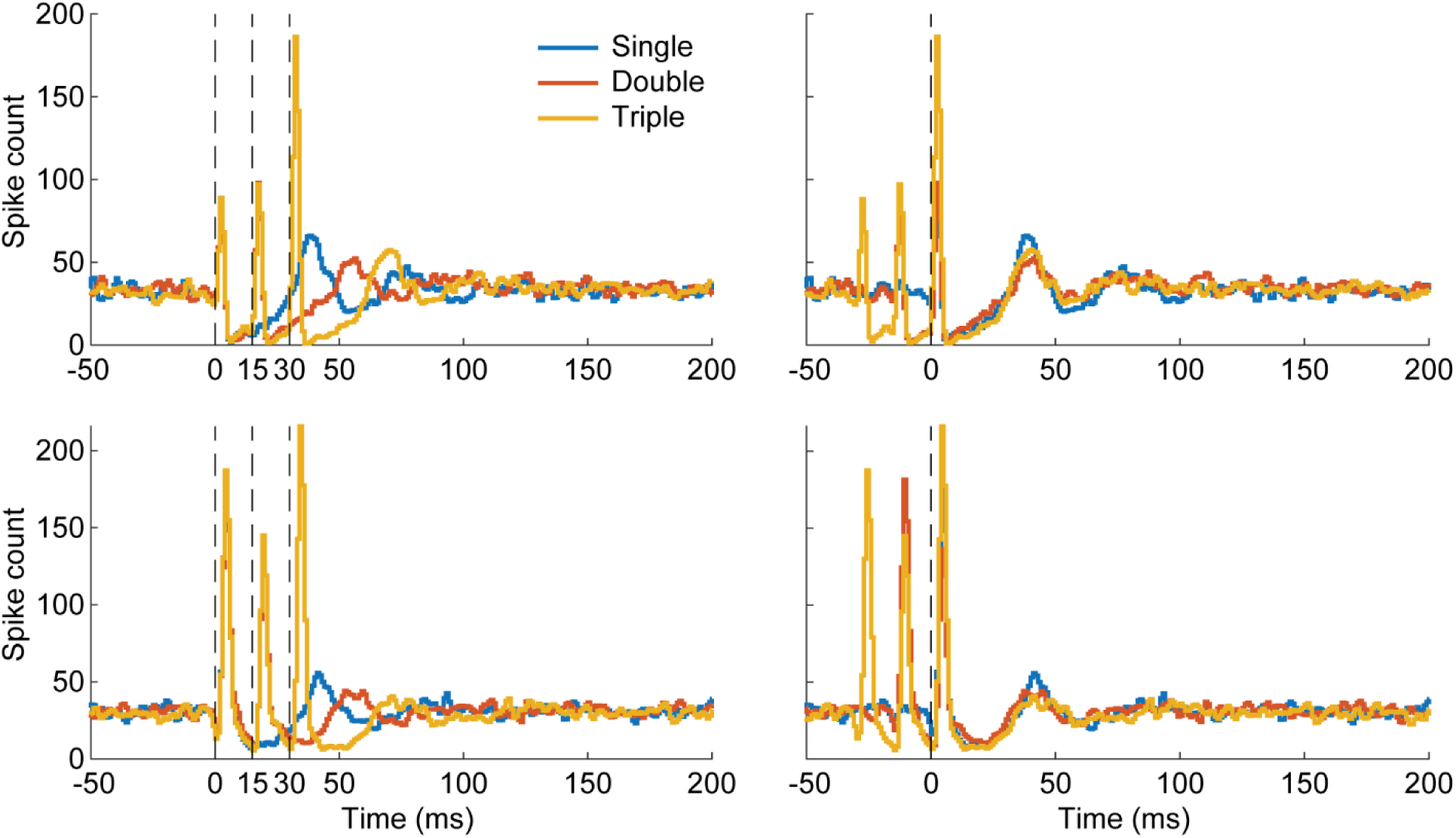
Stimulation during inhibition. Two PSTH examples with 1 ms bins of double- and triple-pulse stimulation in which subsequent pulses arrive during the inhibitory response of the previous pulse (left). Spikes were readily evoked even when stimulating during the inhibitory response. Aligning the PSTHs to the final stimulus pulse (right) shows that the inhibition restarts at each stimulus pulse. Each condition consisted of 1500 stimuli.

**Figure 4.**
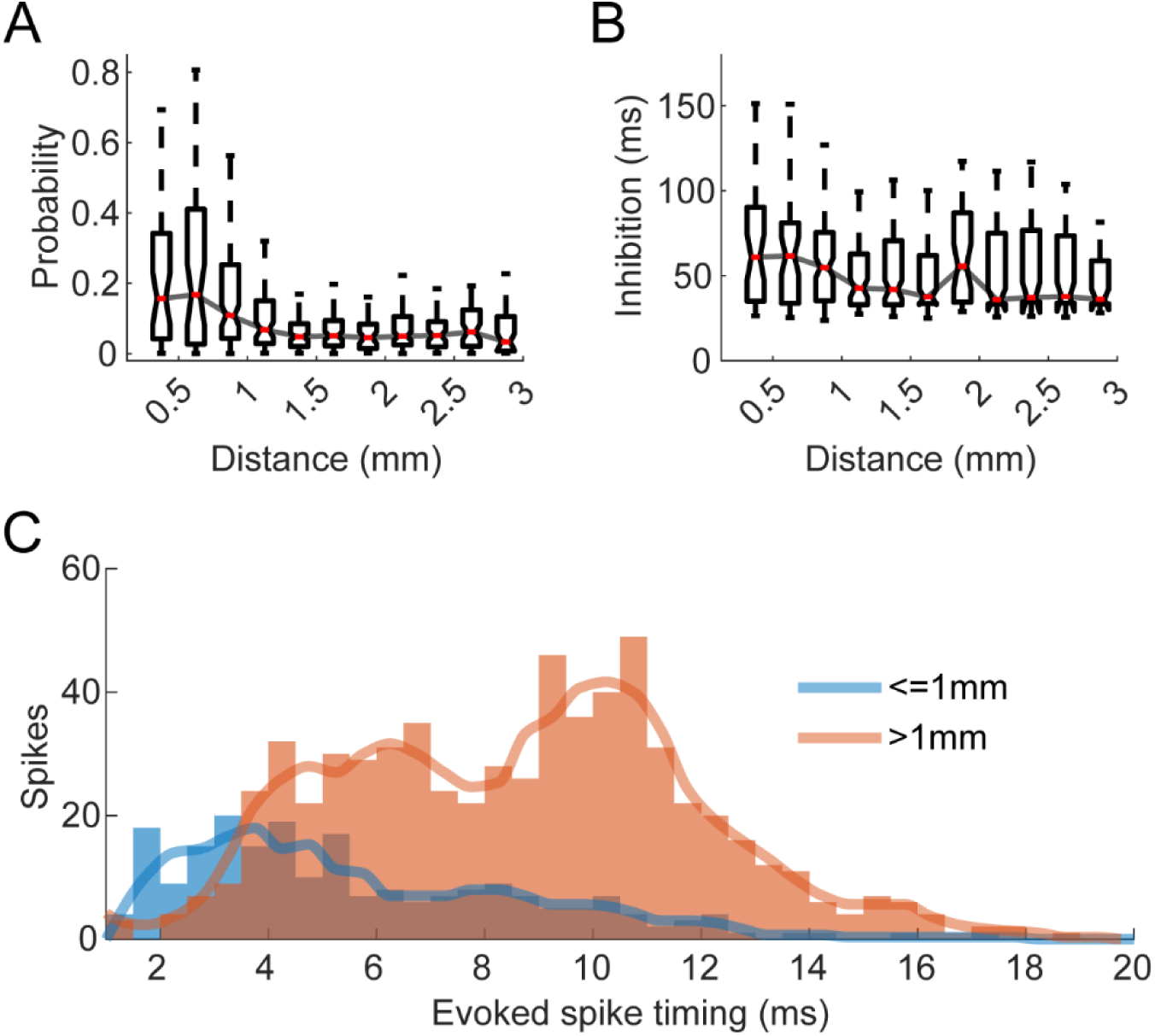
Effect of distance from stimulus site. **A.** Evoked spike probability with respect to distance from the stimulated site. **B.** inhibition duration with respect to distance from the stimulated site. **C.** Histogram of evoked spike timings split into sites close (<1mm) to the stimulated site and all other sites. The line shows the cubic interpolated moving average over 3 bins.

**Figure 5.**
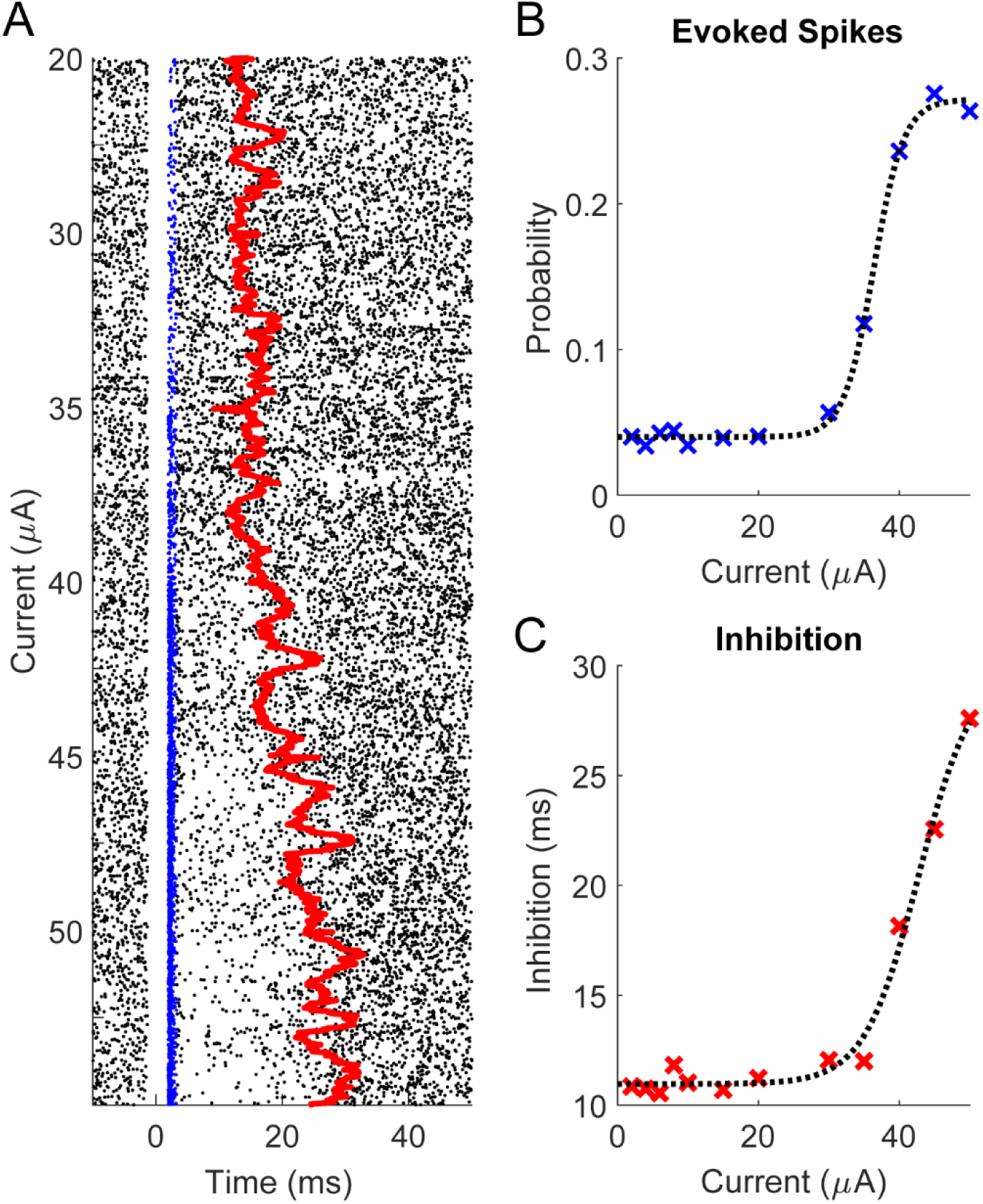
Effect of stimulus amplitude. **A.** An example raster plot of a spike over different stimulus current amplitudes delivered for five minutes each at 2 Hz. Blue dots represent evoked spikes and the red line shows the median of the inhibition duration binned every 30 seconds with 1 second steps. **B.** An example evoked spike probability as a function of amplitude and **c)** inhibition duration as a function of stimulus amplitude. The dashed lines are fitted sigmoidal curves using least-squares regression.

### Effects of distance from stimulation site and stimulus amplitude

Evoked spikes occurred with greater probability and less variable latencies for units close to the stimulus electrode than for more distant units. The probability of evoking a spike in electrodes <1 mm from the stimulus site was significantly greater than for further sites (p=1.3e-14). In addition, closer sites on average had evoked spikes that occurred at shorter, and less variable latencies (p=8.5e-29), suggesting the presence of mono- and polysynaptic activation (Figure 3 – Stimulation during inhibition

Two PSTH examples with 1 ms bins of double- and triple-pulse stimulation in which subsequent pulses arrive during the inhibitory response of the previous pulse (left). Spikes were readily evoked even when stimulating during the inhibitory response. Aligning the PSTHs to the final stimulus pulse (right) shows that the inhibition restarts at each stimulus pulse. Each condition consisted of 1500 stimuli.

**Figure A).**
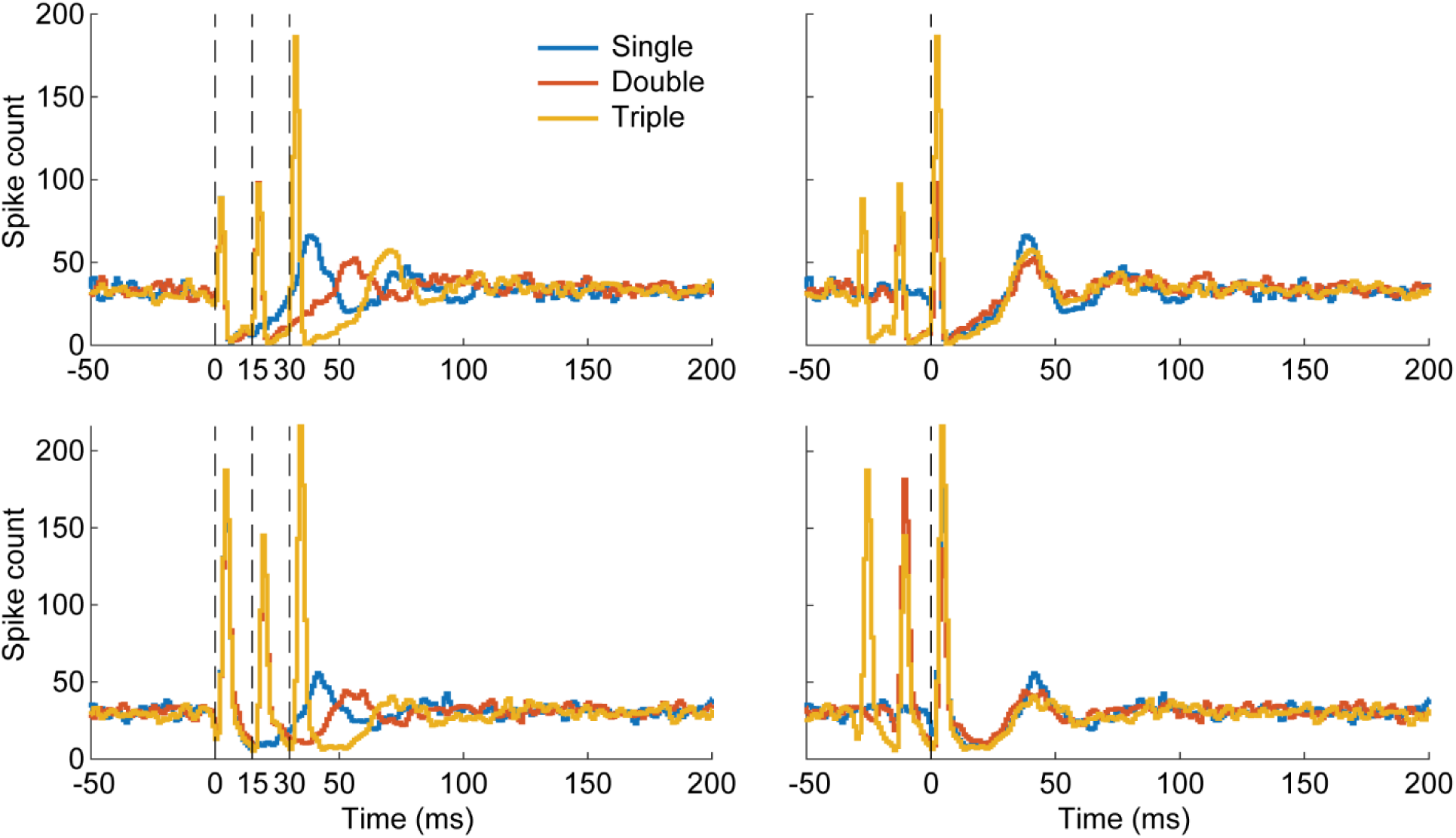
The duration of inhibition had a similar trend: recording sites <1 mm from the stimulus site tended to show stronger inhibition compared to further sites (p=0.003) (Figure 3 – Stimulation during inhibition

Two PSTH examples with 1 ms bins of double- and triple-pulse stimulation in which subsequent pulses arrive during the inhibitory response of the previous pulse (left). Spikes were readily evoked even when stimulating during the inhibitory response. Aligning the PSTHs to the final stimulus pulse (right) shows that the inhibition restarts at each stimulus pulse. Each condition consisted of 1500 stimuli.

**Figure B).**
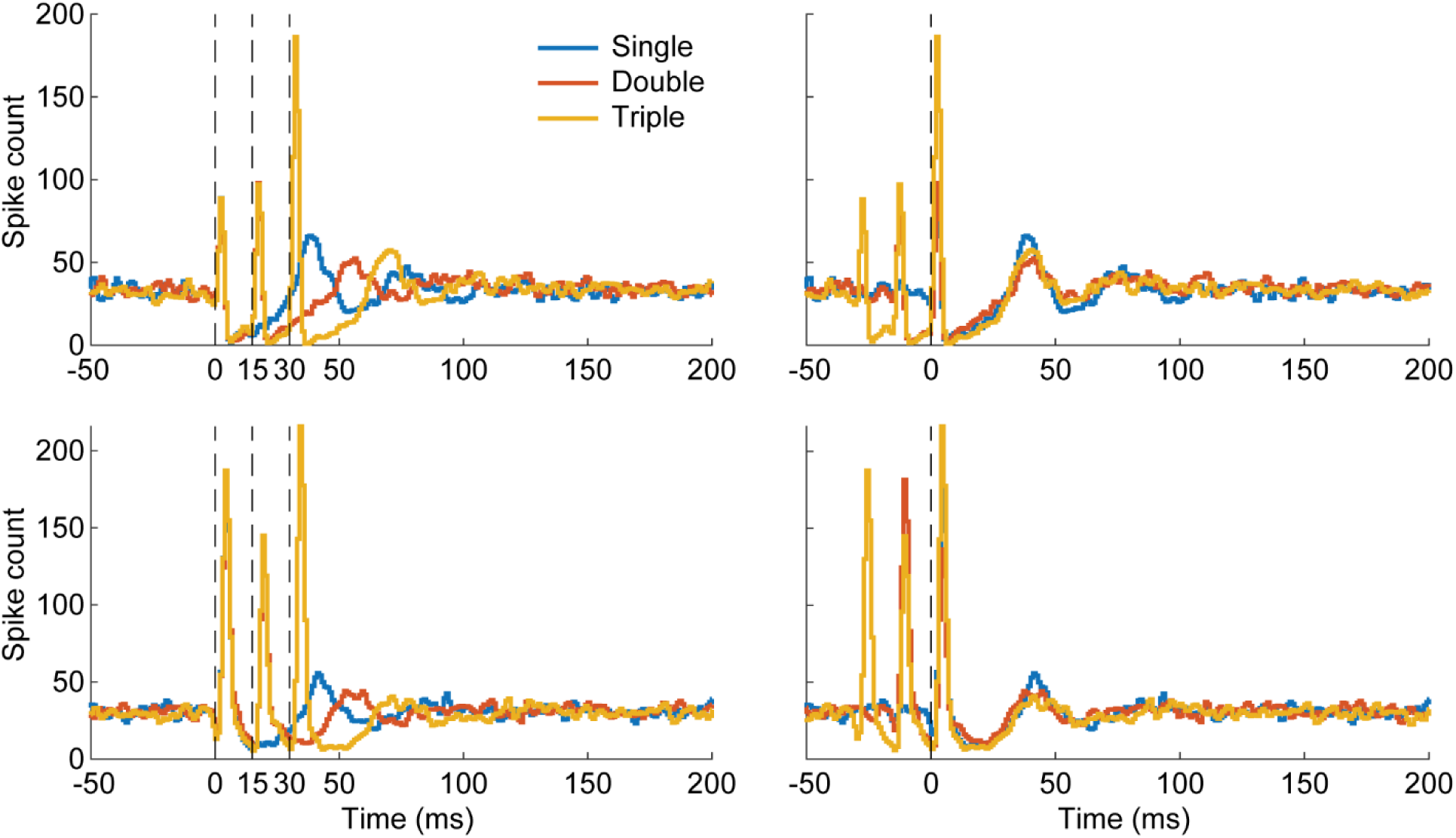
The probability of evoking spikes and the duration of inhibition increased sigmoidally with the stimulus amplitude for all responsive units (Figure). The sigmoid curves were always steep: a change of 10 to 20 μA in stimulus intensity generated the difference between 5% and 95% of the maximum value for both evoked spike probability and inhibition duration.

### Covarying evoked spikes

If stimulus evoked spikes are activated by cortical circuitry, there must be covarying responses between different spikes. Thus, we determined whether pairs of units were likely to have the same responses for individual stimuli. Figure 6A shows the Pearson correlation coefficient between pairs of units plotted against distance from one another. There are clearly pairs with significant correlations, though the correlations are usually weak with the vast majority of correlations having a correlation coefficient less than 0.2. Pairs of units closer together have a significantly higher chance to be correlated, as they are likely modulated by similar circuitry (Figure 6A, right).

**Figure 6.**
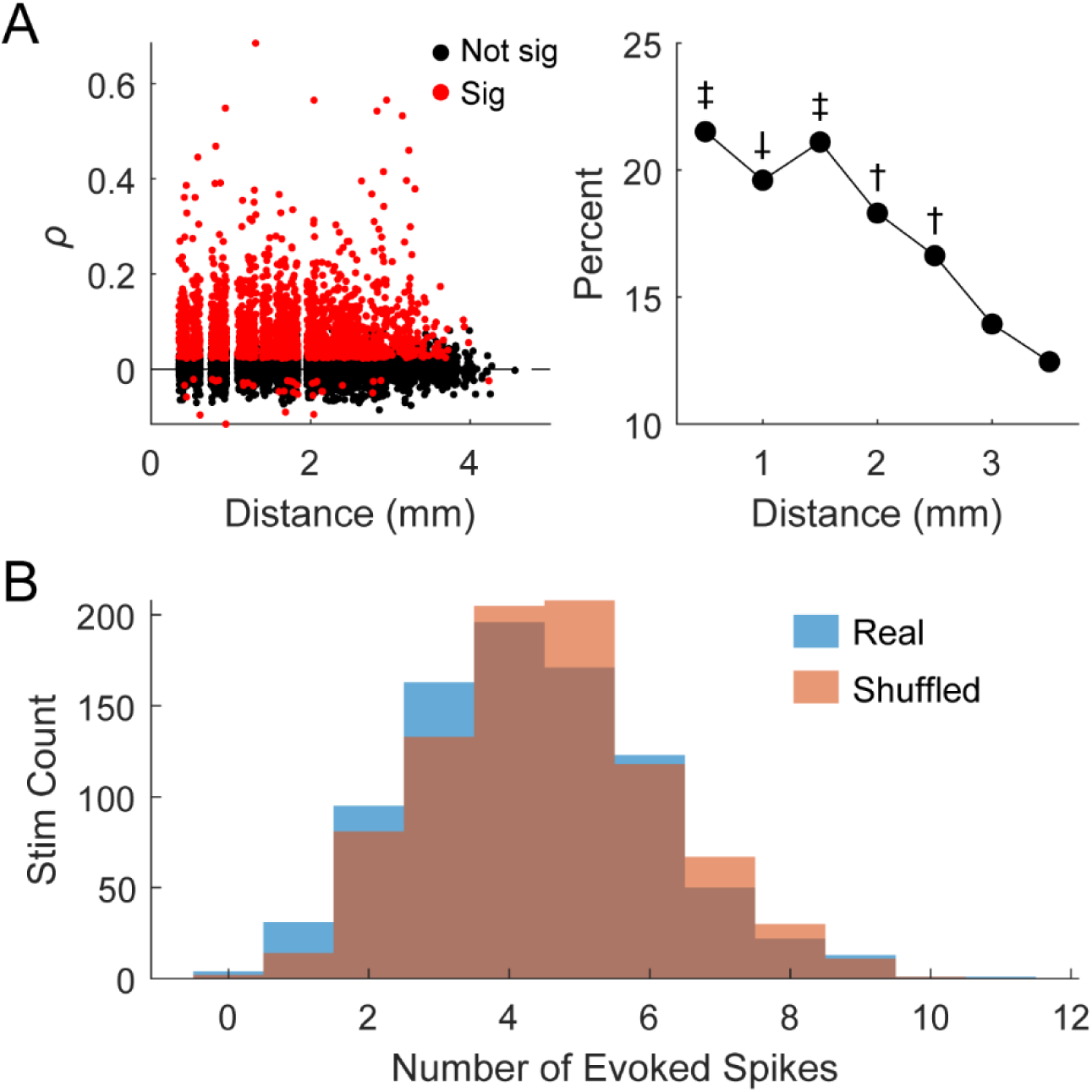
Covarying evoked spikes. **A.** Pearson correlation coefficient between pairs of units on whether individual stimuli evoked a spike or not plotted against the distance between the units (left). Pairs of units closer to each other had a higher chance of having a significant correlation (right; †: significantly different from the last two points; ‡: significantly different from the last 4 points; ⸸: significantly different from the last 3 points; p<0.05, ANOVA). **B.** An example of distributions of stimuli that evoked a specific number of evoked spikes using the true data (True) and when the evoked spikes were shuffled (Shuffled). The two distributions were not statistically significant from one another (p=0.11, Kolmogorov-Smirnov test).

The presence of covarying pairs at large distances (>3 mm) led us to analyze whether there was a population wide response driven by an underlying cortical state. We first collected the total number of evoked spikes generated by each stimulus. Then for each evoked spike we shuffled which stimulus generated a spike and recalculated the total number of evoked spikes attributed to each stimulus. The existence of population dependence should show that the distributions of the original counts and the shuffled counts should be significantly different. However, we found that the two distributions are not significantly different to one another in any session (Figure 6B).

### Spontaneous activity affects stimulus responses

In addition to stimulation current and separation of the recording and stimulation sites, we found two other dependencies that affect the response. One is the spontaneous firing rate of the recorded units. Both the evoked spike probability and inhibition duration had statistically significant correlations with spontaneous firing rate over time (Figure 7). Table 1 documents the number of units with uncorrelated, positively correlated, or negatively correlated evoked spike probability and inhibition duration with spontaneous firing rate. A slight majority of units tested (300/585, 51%) had evoked spike probabilities that were positively correlated with firing rate and inhibition duration that were negatively correlated with firing rate. We additionally performed a 2-way ANOVA to determine whether the two relationships were dependent on one another but found no significant relationship (p=0.11). Units with positively correlated evoked spike probabilities and spontaneous firing rate were typically farther from the stimulated site than units with a negatively correlated relationship (Figure 7B left, 7C left). In contrast, units with positively correlated inhibition duration and spontaneous firing rate were typically closer to the stimulated site (Figure 7B right, 7C right).

**Figure 7.**
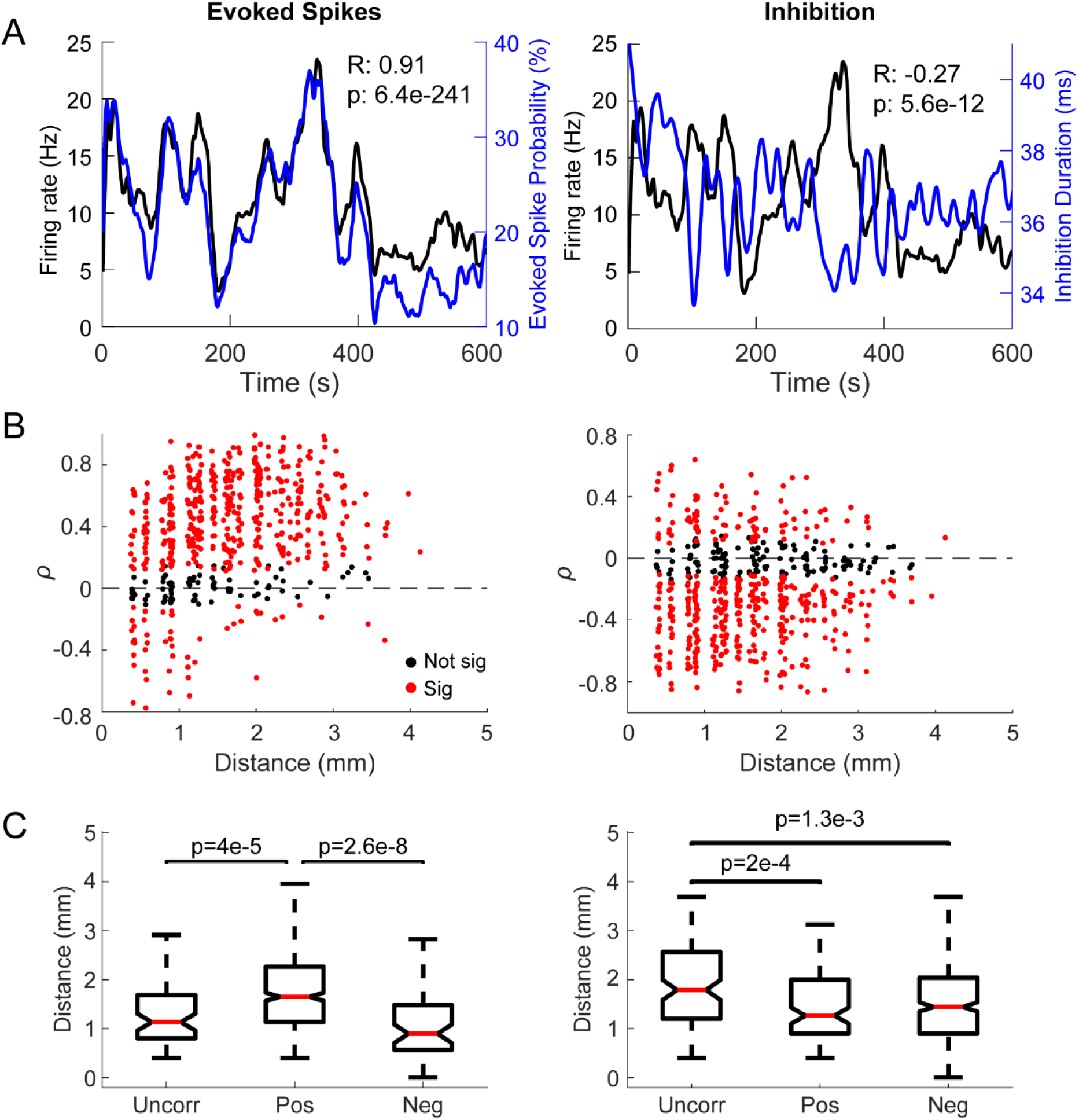
Probability of evoking a spike and inhibition duration are related to spontaneous firing rate. **A.** An example of a neuron with positively correlated firing rate (black) and evoked spike probability (blue) over 10 minutes. The rate and probabilities are averaged over 30 second bins with 1 second steps. **B.** Scatter plot of the Pearson correlation coefficient (*ρ*) between the spontaneous firing rate and the probability of evoking spikes (left) or the inhibition duration (right) against the distance of the recorded spike from the stimulated site. **C.** The same unit had a negatively correlated firing rate (black) and inhibition duration (blue). The inhibition duration is also averaged over 30 second bins with 1 second steps. **C.** Distance from the stimulated site for units with uncorrelated, positively correlated, and negatively correlated evoked spike probabilities (left) or inhibition duration (right) with firing rate. Labeled p-values are from the Wilcoxon rank-sum test.

**Table 1.** Evoked spike probability and inhibition duration correlations with spontaneous firing rate.

|  |  | Inhibition duration and spontaneous firing rate |  |  |  |
| --- | --- | --- | --- | --- | --- |
|  |  | Uncorrelated | Positive<br>Correlation | Negative<br>Correlation | Total |
| Evoked spike<br>probability and<br>spontaneous<br>firing rate | Uncorrelated | 10 | 18 | 37 | 65 |
|  | Positive<br>Correlation | 86 | 77 | 300 | 463 |
|  | Negative<br>Correlation | 12 | 13 | 32 | 57 |
|  | Total | 108 | 108 | 369 | 585 |

The second dependency was the timing of stimuli relative to the most recent spike. We analyzed 17 units (2 from monkey S, 3 from monkey K, and 12 from monkey J). In 14 of the 17 units (2 from monkey S, 3 from monkey K, and 9 from monkey J), the probability of evoking a spike varied as a function of the time between the onset of stimulation and the most recent spontaneous spike. For these 14 units the probability was significantly positively correlated with the unit’s autocorrelogram in the absence of stimulation (Figure 8, left and middle). The probability distributions were the same even if the most recent spike was evoked by stimulation rather than being spontaneous. The inhibitory response did not depend on the timing of pre-stimulus spikes.

**Figure 8.**
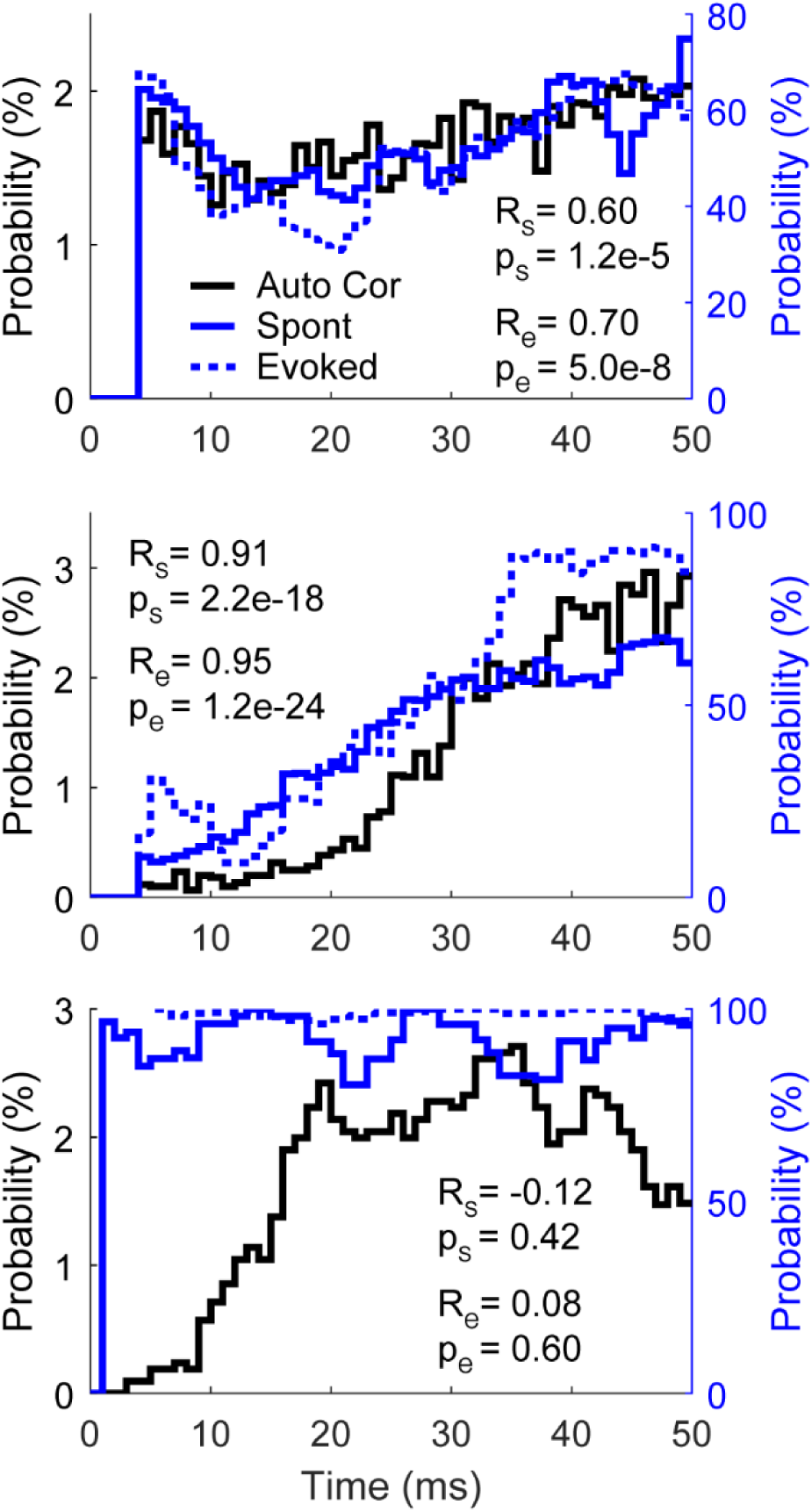
Probability of evoking a spike is dependent on the timing of stimulus. Three examples of spike autocorrelations (Auto Cor), and probability of a stimulus evoking a spike relative to timing from the most recent spontaneous (Spont) and evoked (Evoked) spike. Top and middle show two different autocorrelation waveforms with correlated evoked spike probability. They are not aligned in the bottom example; this typically occurs when the probability of evoking a spike is high. All traces show a moving average using 5 ms bins with a 1 ms step size. R_s_ is the correlation coefficient between Auto Cor and Spont, R_e_ the correlation coefficient between Auto Cor and Evoked, and p_s_ and p_e_ are the corresponding p-values of correlation.

In 3 of the 17 tested units the probability of evoking a spike did not depend on the timing between the previous spike and stimulus onset. In all 3 of these units the probability of evoking a spike was consistently high (Figure 8, right), which suggests that the stimulation amplitude was large enough to evoke spikes regardless of other properties.

### Repetitive stimulation changes stimulus response over time

Repetitive microsimulation has been shown to modulate the responses of units to ICMS (Dadarlat et al., 2019; Michelson et al., 2019; Zucker & Regehr, 2002). We documented the effects of repetitive ICMS over the stimulus period ranging from 5 to 20 minutes on the evoked spike probability and the duration of inhibition by delivering stimuli at 2, 5, 10, or 20 Hz. The probability of evoking a spike often increased or decreased over time, following a linear or exponential trend over the course of the session (Figure 9A).

**Figure 9.**
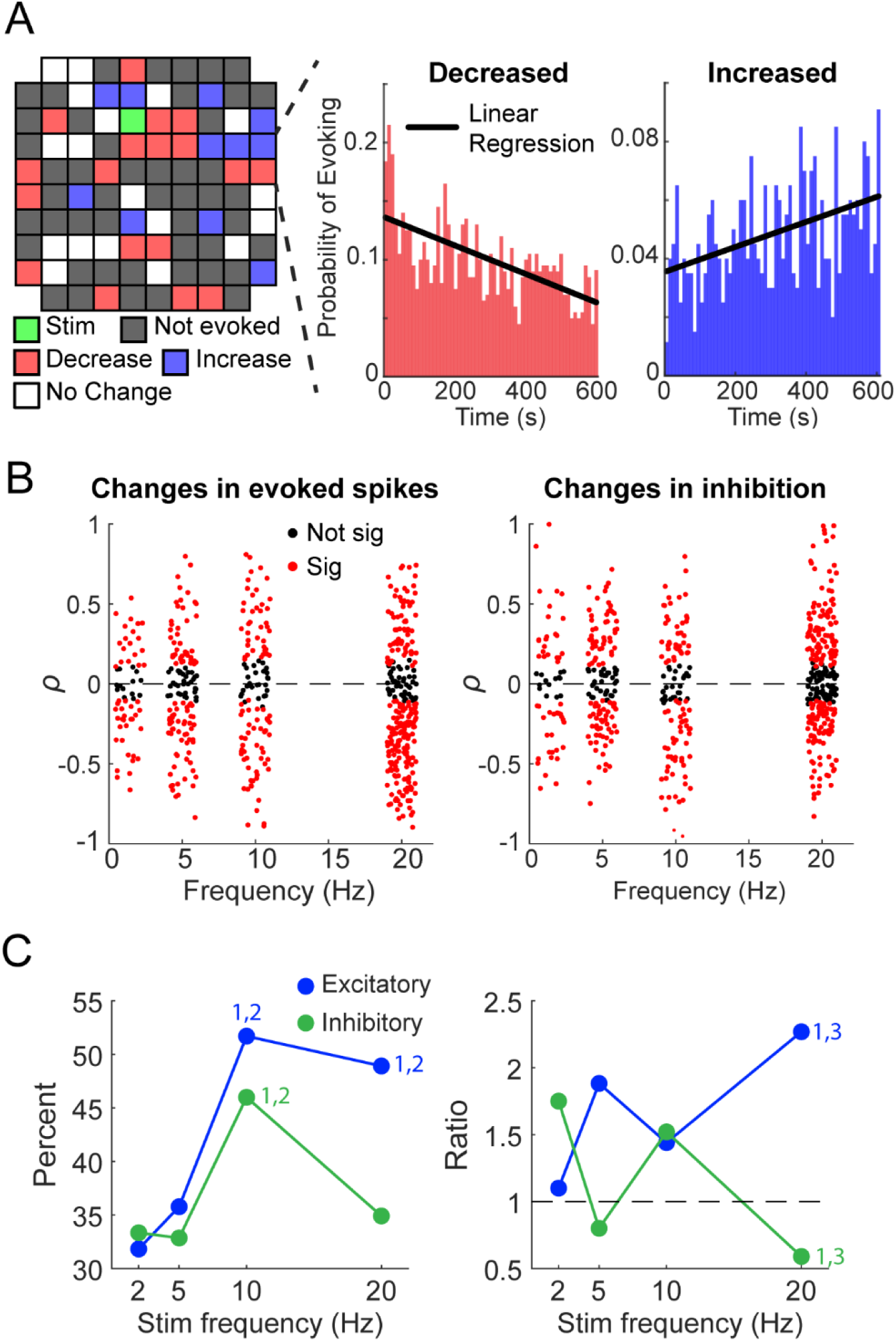
Changes in evoked spike probability and inhibition duration with repetitive stimulation. **A.** Left: Changes in evoked spikes across the array during a session with 10 Hz repetitive stimulation. A random unit was chosen for each electrode to demonstrate the lack of spatial organization of changes in responses. Right: Examples of changes in the probability of evoking spikes increasing or decreasing over time. **B.** Pearson correlation coefficients of evoked spike probability and inhibition duration plotted against stimulation frequency. Note that experiments delivered tonic stimulation at 2, 5, 10, or 20 Hz; a jitter was added to the frequencies of each point to better visualize the data. **C.** The percentage of spikes that had significant (p<0.05) changes over time for each stimulation frequency for both evoked spike probability and inhibition (left) as well as the ratio of decreases to increases (right). The dashed line of the right plot shows a threshold – if the value is higher (>1) the changes induced are more likely to be decreasing whereas if the value is lower (<1) the changes are more likely to be increasing. Numbers above points denote significance (1: significant from 2 Hz, 2: significant from 5 Hz, 3: significant from 10 Hz; p<0.05, ANOVA).

The evoked spike probability was significantly more likely to change with higher frequencies of stimulation (Figure 9B left, 9C left). Of the changes, high frequencies (20 Hz) were likely to cause decreases in evoke spike probability compared to lower frequencies. Changes in inhibition duration over time were also more likely to occur at higher frequencies, and high frequencies were more likely to cause increases in inhibition duration. (Figure 9B right, 9C right)

The changes also depended on the distance from the recording site to the stimulated site (Figure 10A). The probability of evoking spikes and inhibition duration were significantly more likely to change the closer the units were to the stimulated site (Figure 10B, left). The specific change in evoked spike probability did not have a dependence on distance, but inhibition duration was more likely to decrease in sites further from the stimulated site (Figure 10B, right).

**Figure 10.**
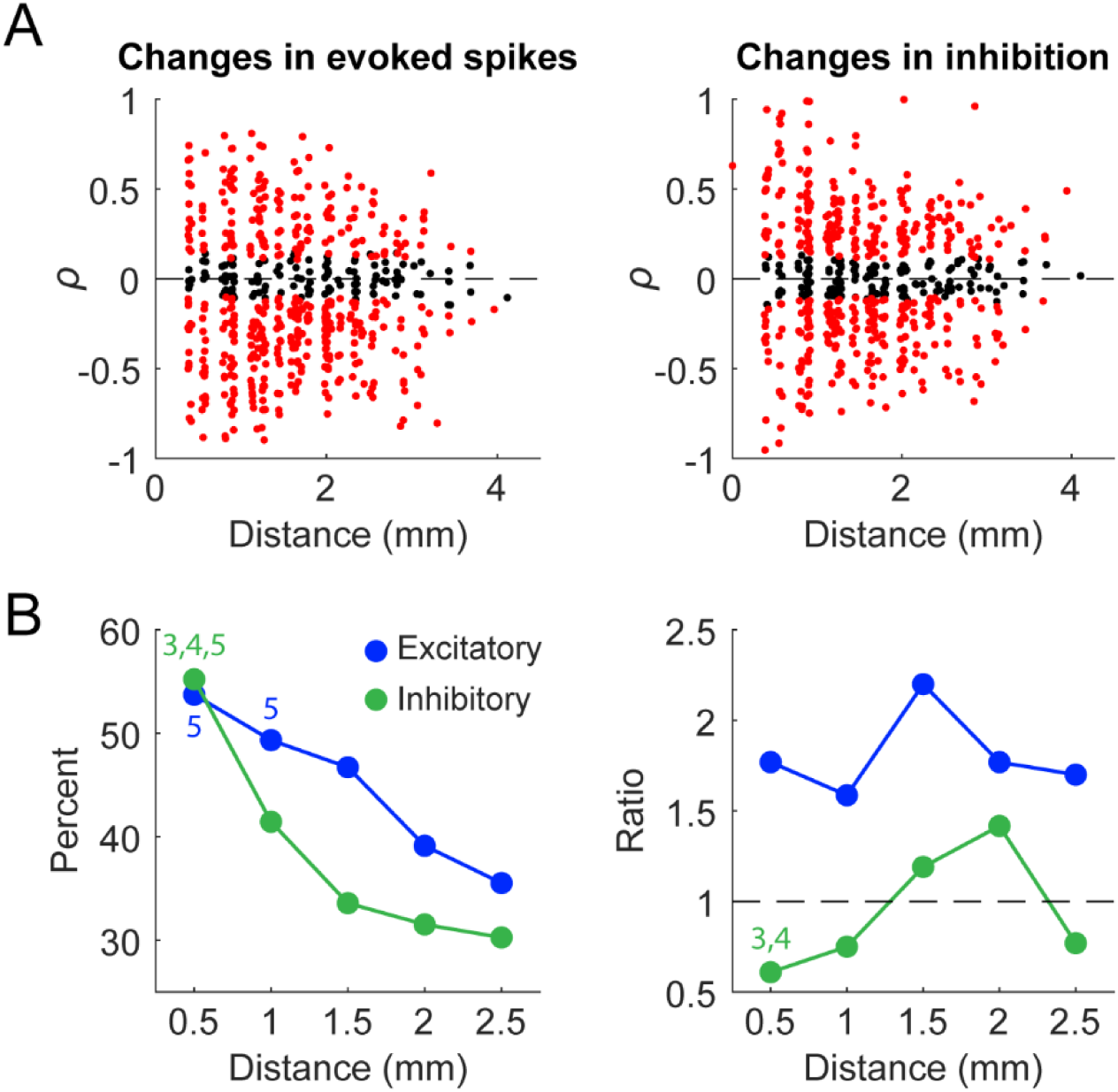
Changes with repetitive stimulation with respect to distance. **A.** Pearson correlation coefficients of evoked spike probability and inhibition duration plotted against distance of the recorded spike from the stimulated site. Note that experiments delivered tonic stimulation at 2, 5, 10, or 20 Hz; a jitter was added to the frequencies of each point to better visualize the data. **B.** The percentage of spikes that had significant (p<0.05) changes over time for each bin of distance (±0.25 mm around each point) for both evoked spike probability and inhibition (left) as well as the ratio of decreases to increases (right). We did not include data points from channels greater than 2.75 mm from the stimulated site to this figure due to the lack of samples. Numbers above points denote significance (3: significant from 1.5 mm, 4: significant from 2 mm, 5: significant from 2.5 mm; p<0.05, ANOVA).

Finally, in 72/585 (12%) of units, we also observed changes over time in the latency of the evoked spikes (Figure 11A). Latency changes typically occurred in units that were recorded on electrodes closer to the stimulation site, more commonly occurred with high frequency stimulation, and was more likely to increase over time (Figure 11B). Of the units with changes in their evoked spike latency, 249/447 (56%) units had increasing latencies and 198/447 (44%) had decreasing latencies. Distance from the stimulated site also played a role, with units closer to stimulation more likely to have changes in their evoked spike latency over time (Figure 11C).

**Figure 11.**
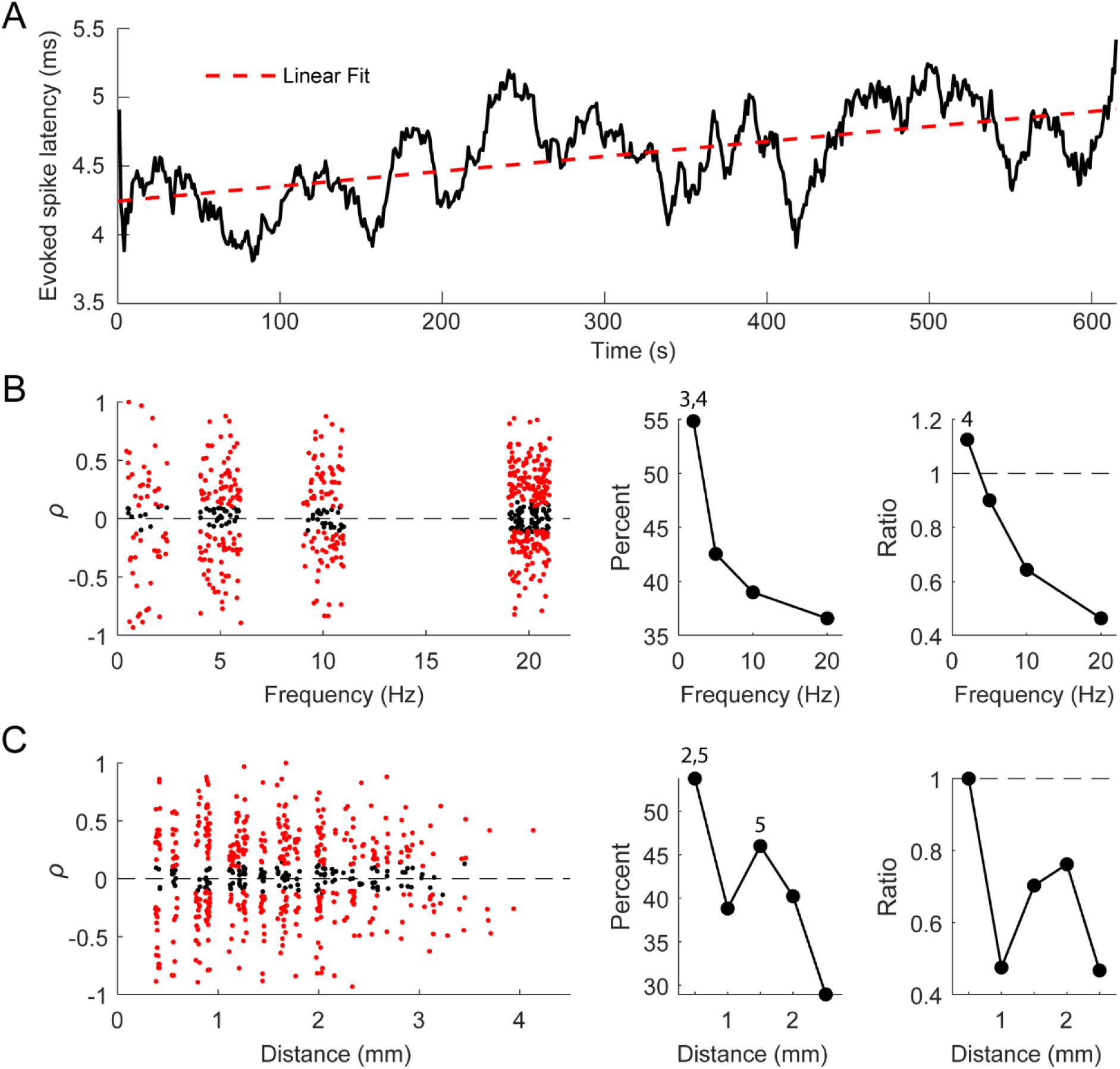
Changes in evoked spike latency with repetitive stimulation. **A.** An example of evoked spike latency changing over time. **B.** Scatter plot of Pearson correlation of the evoked spike latency over time with respect to stimulation frequency (left). Percentage of evoked spike latencies with a significant change over time (middle), and the ratio of decreases to increases (right). Numbers above points denote significance (3: significant from 10 Hz, 4: significant from 20 Hz; p<0.05, ANOVA). **C.** Scatter plot of Pearson correlation of the evoked spike latency over time with respect to distance from the stimulated site (left). Percentage of evoked spike latencies with a significant change over time (middle), and the ratio of decreases to increases (right). Numbers above points denote significance (2: significant from 1 mm, 5: significant from 2.5 mm; p<0.05, ANOVA).

### Relationship between evoked spikes and inhibition

Since evoked spikes and inhibition often both exhibited changes over time, we sought to determine whether they were directly related on a trial-by-trial basis. Individual stimuli in each experiment were divided into two categories: those that evoked spikes and those that did not. For the unit in Figure 12A the PSTHs of the two classifications of stimuli are very similar except for the evoked spike peak. We found no statistical pairwise difference between the inhibition induced by stimuli that evoked spikes compared to the stimuli that did not evoke spikes (585 units, Figure 12B).

**Figure 12.**
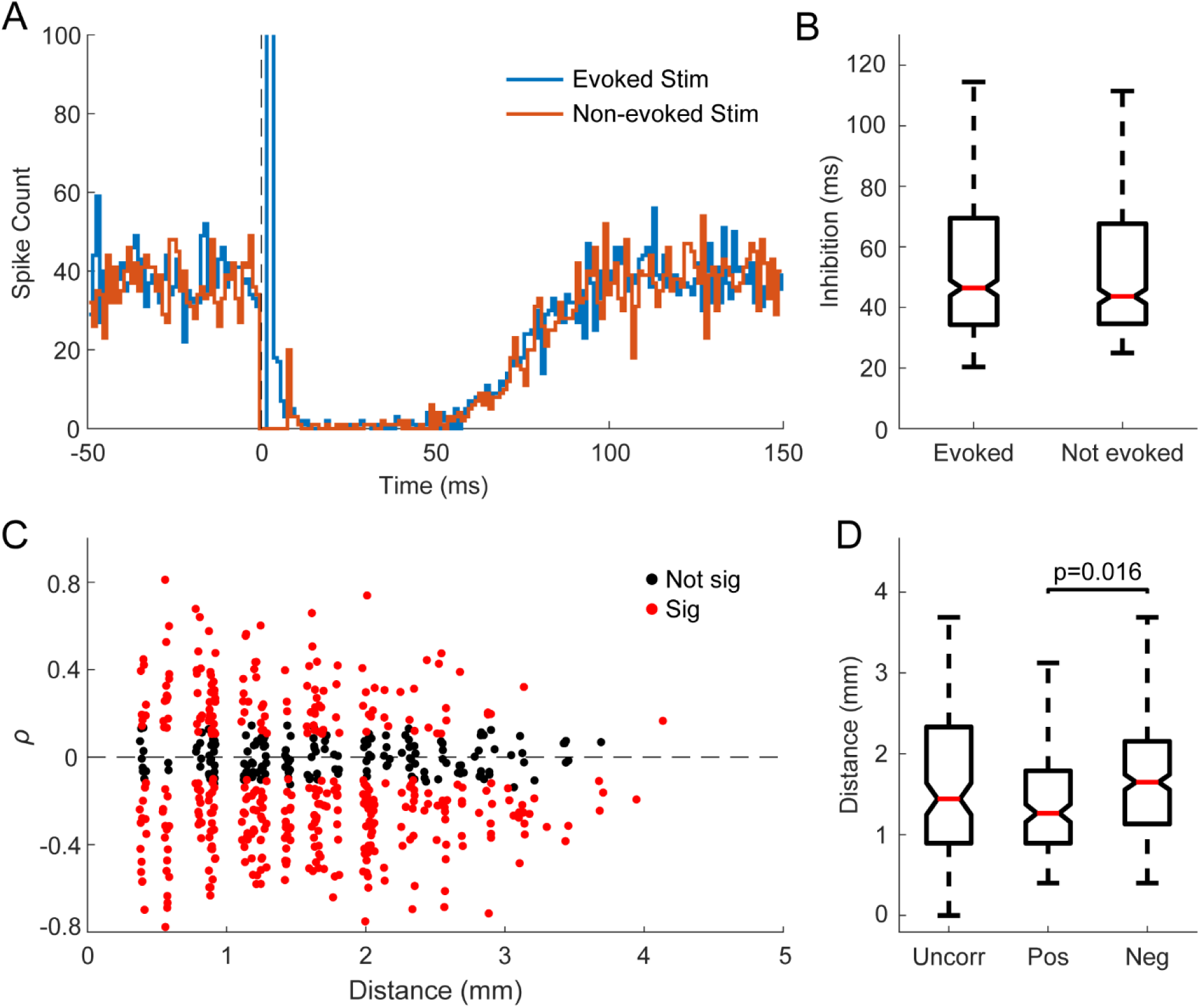
Relationship between evoked spikes and inhibition. **A.** Example PSTH with 1 ms bins following stimuli that evoked spikes and those that did not demonstrating similar inhibitory response. Each condition consisted of 1500 stimuli. Smoothed (2 ms wide gaussian moving window) PSTH of the two different stimulation classifications. Note the inhibition is extremely similar for both. **B.** A comparison of the inhibition strength in the two different classifications for 470 units. There was no statistically significant pairwise difference between the two groups. **C.** Scatter plot of the Pearson correlation coefficient between evoked spike probability and inhibition duration against distance of the recorded spike from the stimulated site. **D.** Comparisons of distance from the stimulus site of units with uncorrelated, positively correlated, and negatively correlated evoked spike probability and inhibition duration. The labeled p-value is from the Wilcoxon rank-sum test.

Although whether a stimulus evokes a spike does not affect the inhibitory response, the mechanisms underlying the changes over time could nevertheless be related. We analyzed the correlation between the probability of evoking a spike and duration of inhibition over time to determine whether they were positively or negatively correlated. Correlations with p<0.05 were considered significant, and all other instances were denoted to be uncorrelated. Of the 585 units tested, 30% (173 units) had positively correlated changes in the probability of evoking spikes and the duration of inhibition, 47% (273 units) had negatively correlated changes, and 23% (135 units) were uncorrelated. Units with positively correlated evoked spike probabilities and inhibition duration tended to be closer to the stimulated site (Figure 12C and 12D).

### Spike type does not correlate with stimulus response properties

To determine whether the cell type of the recorded units influenced their responses to ICMS, we used the spike width, calculated as the time between the minimum value of the waveform to the maximum value, to classify each neuron as fast-spiking (FS) or regular-spiking (RS) (Connors & Gutnick, 1990; McCormick et al., 1985).

Figures 13A shows an example of the two different spike waveforms, and Figure 13B shows the distribution of their spike widths. We separated the units into two groups based on their defined spike width shown at the dotted line in Figure 13B. Around 10% of all units fell to the left of the line and were denoted to be FS; the rest were denoted to be RS. We found that the putative cell type did not correlate with the distribution of distances from the stimulated site, whether a spike could be evoked, the probability of evoking a spike, the duration of inhibition, or how any measure changed over time (Figures 13C-G). Thus, all results reported herein are independent of the type of neuron recorded.

**Figure 13.**
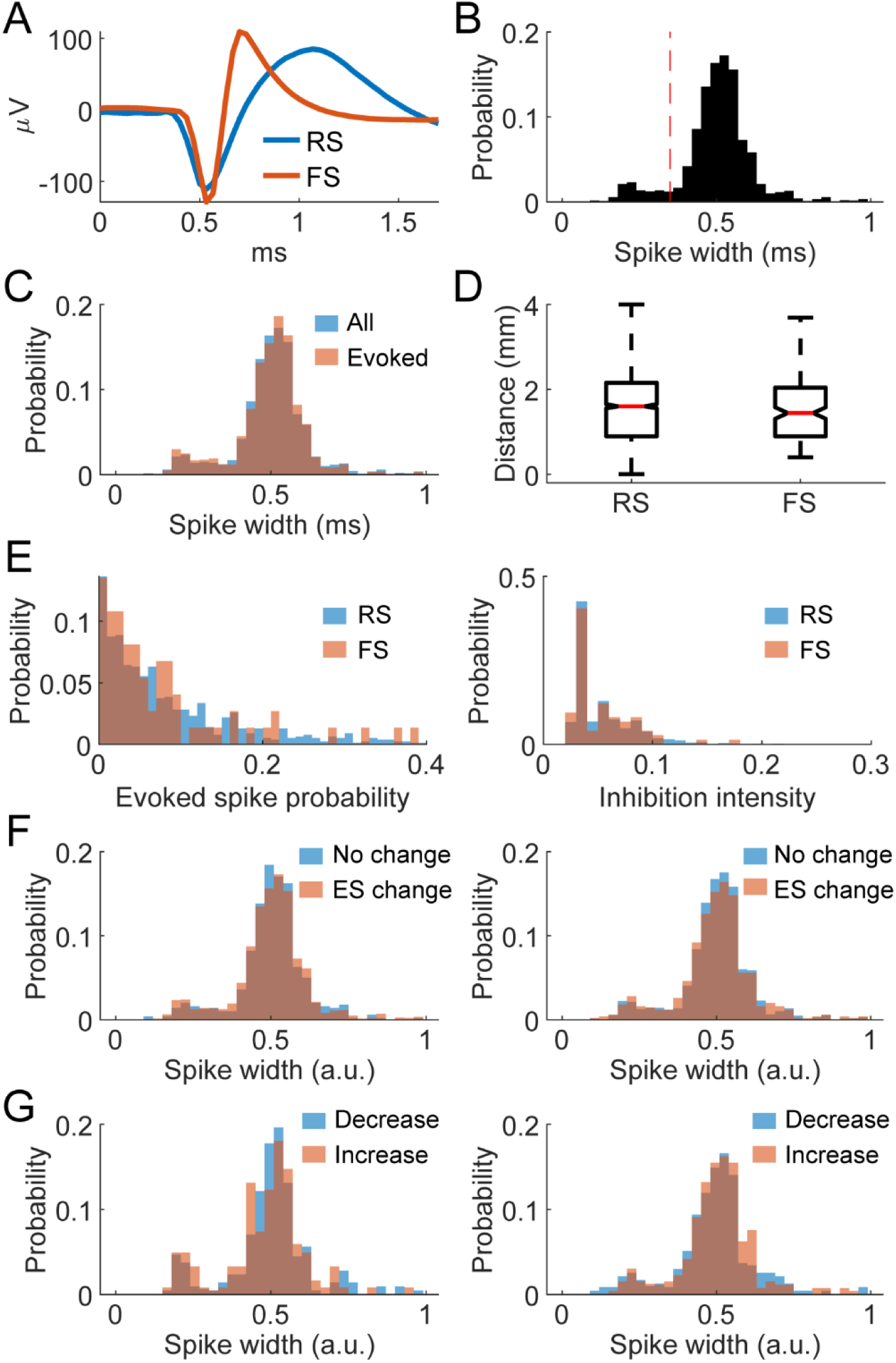
Comparisons between regular spiking and fast spiking neurons. **A.** Example of regular spiking (RS) and fast spiking (FS) neuron waveforms. **B.** Distribution of the widths of spike waveforms (trough to peak time). Vertical dotted line indicates the classification boundary (0.35 ms) – fast spiking neurons fall to the left and regular spiking to the right. **C.** Spike width distribution of all recorded spikes and spikes that were evoked by stimuli. **D.** Distance from the stimulated site of evoked spikes grouped by spike width. **E.** Evoked spike probability distribution (left) and inhibition duration distribution (right) of RS and FS neurons. **F.** Spike width distributions of evoked spike probability change vs no change over time (left) and inhibition duration change vs no change over time (right). **G.** Spike width distributions of evoked spike probability decrease vs increase over time (left) and inhibition duration decrease vs increase over time (right).

## Discussion

### Comparisons to previous studies

ICMS has been shown to predominantly activate neurons transsynaptically (Butovas & Schwarz, 2003; Hussin et al., 2015; Klink et al., 2017). We observed in our study that spike latencies fluctuated more than would occur with antidromic activation, and stimuli that were delivered within 1 millisecond after a spike were still able to evoke spikes. Since antidromic activation would result in collision and an absence of a spike at such short latencies, this suggests that the recorded units were predominantly activated orthodromically. Also consistent with previous experiments, we saw evoked spikes in units recorded up to 4.5 mm away from the stimulation site, which suggests that ICMS activates a distributed population rather than only a concentrated sphere of neurons around the electrode tip (Butovas & Schwarz, 2003; Hao et al., 2016; Histed et al., 2009).

Although the excitatory response was directly measurable in our experiments, the subsequent inhibitory response manifested as a lack of spikes. Butovas et al. concluded that a similar inhibitory response to ICMS was likely caused by GABA_B_ receptors, which they confirmed with a follow-up study with pharmacological blocks (Butovas et al., 2006; Butovas & Schwarz, 2003). GABAergic inhibition would also explain the similarity in the sigmoidal curves of evoked spikes and inhibition with stimulus amplitude in our experiments – if more excitatory neurons are excited by the higher intensity stimulation, more inhibitory neurons will be activated via feedforward and feedback circuits, thereby increasing the amount of inhibition (Figure 14). A lack of relationship between inhibition and distance to stimulation site as well as the presence of inhibition in units without evoked spikes is likely due to the high connectivity of interneurons compared to principal cells (Isaacson & Scanziani, 2011; Matsumura et al., 1996; Rudy et al., 2013; Tremblay et al., 2016). Interestingly, Hao et al. found that inhibition decreased as a function of distance, which we did not find (Hao et al., 2016). Potential reasons for this discrepancy could be the differences in how we measured inhibition and the stimulus amplitude.

**Figure 14.**
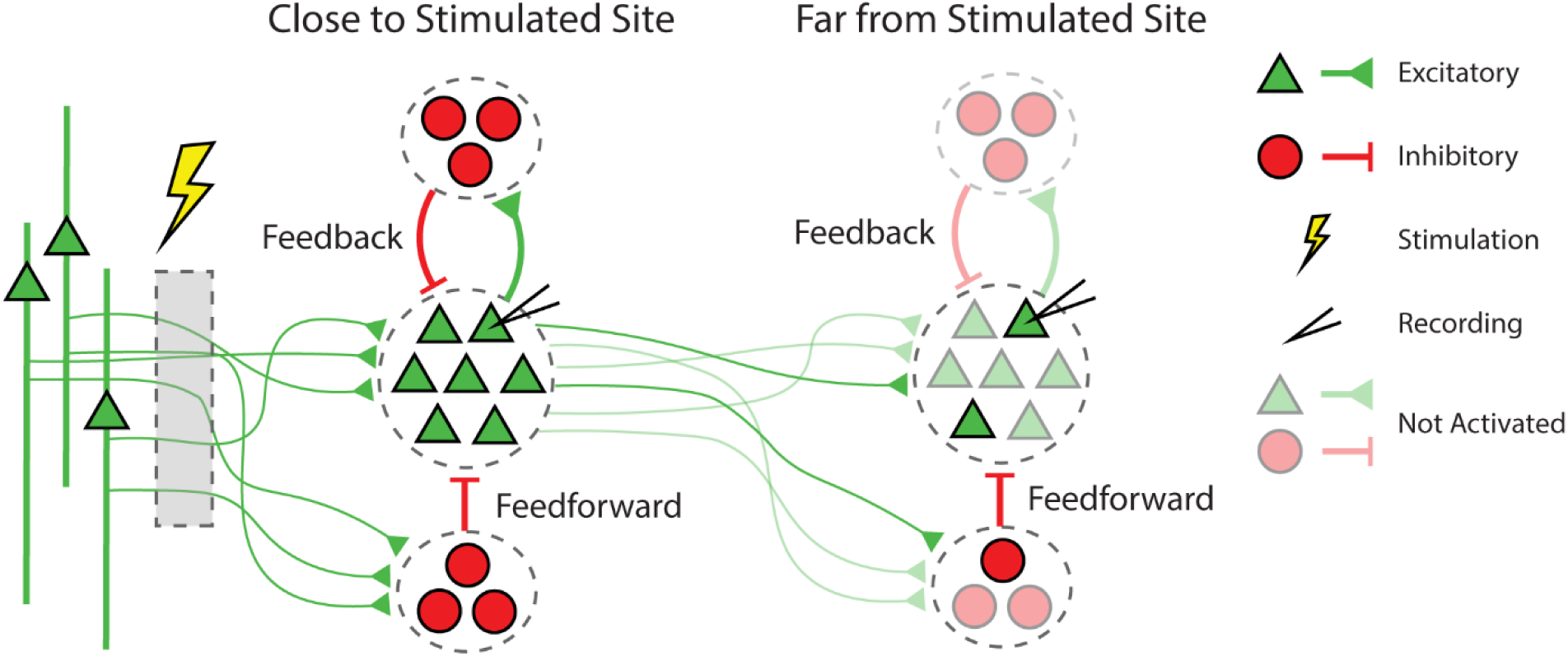
Stimulation response schematic. Schematic of ICMS activated circuitry that generates excitatory and inhibitory responses through feedforward and feedback mechanisms. Stimulation activates axons projecting to the recording site. Sites closer to the stimulated site have more complete activation compared to sites further from the stimulated site. Possible direct connections from the stimulated site to the far site would also be sparsely activated.

However, we found that inhibition typically lasted between 5-100 ms and rarely over 100 ms, which is significantly shorter than the average time constants of GABA_B_ inhibitory postsynaptic potentials of 150-200 ms (Bettler et al., 2004; Connors et al., 1988). Therefore, GABA_A_ mediated inhibition is a more likely candidate to explain our results; previous studies have shown that GABA_A_ is involved in recurrent polysynaptic inhibition, which we are likely activating via ICMS (Silberberg & Markram, 2007; Zhu et al., 2011). The different animal models and recorded cortical region in these studies may account for these discrepancies.

ICMS activates long horizontal fibers to both feedforward inhibitory networks and the recorded excitatory neurons (Figure 14). These afferent fibers have stronger excitatory connections to the inhibitory interneurons than the principal cells, particularly in layer 2/3, which may explain why we sometimes observe an inhibitory response without any excitatory response (Adesnik & Scanziani, 2010; Cruikshank et al., 2007; Helmstaedter et al., 2008). This, coupled with the fact that inhibitory neurons often target somatic or perisomatic compartments (Isaacson & Scanziani, 2011; Rudy et al., 2013; Tremblay et al., 2016), suggests that the observed inhibition may be initially activated via feedforward circuitry, and is subsequently followed by the excitatory response. The excited principal cells may then activate feedback circuitry that contributes to the inhibitory response (Figure 14).

### Stimulus response depends on network activity and intrinsic membrane properties

Previous research has shown that stimulus responses depend on network activity both *in vitro* (Kumar et al., 2016; Weihberger et al., 2013) and *in vivo* (Kara et al., 2002). Our findings were consistent with these results; the probability of evoking a spike was often positively correlated with spontaneous firing rate, whereas the duration of inhibition was often negatively correlated with firing rate.

English et al. demonstrated that the transmission probability for postsynaptic spikes of inhibitory neurons in the hippocampus *in vivo* is a function of the timing between the previous postsynaptic spike and presynaptic spike (English et al., 2017). Moreover, that study found that this dependency was independent of whether the previous postsynaptic spike was spontaneous or evoked, which suggested that intrinsic properties of the postsynaptic membrane were responsible for the dependence. Our results extend these outcomes by demonstrating that the timing of the previous spike affects not only the transmission probability for spontaneous presynaptic spikes, but also the stimulus-evoked spikes in cortical neurons. Furthermore, we found that the probability distributions for evoking a spike as a function of the timing between the previous spike time and stimulation onset and the spike train autocorrelation were often significantly positively correlated, which further reinforces the notion that this dependency reflects the intrinsic properties of the recorded units.

Altogether, our results reveal that the stimulus response has at least two dependencies other than stimulus amplitude and distance from stimulus site: the intrinsic membrane properties of the recorded neurons and the activity of the network. However, a sufficiently large stimulus current may saturate the responses and overcome these dependencies.

### Repetitive stimulation modulates stimulus responses

Michelson et al. showed that the number of neurons activated by electrical stimulation diminished over time with higher frequencies measured with calcium imaging in a 407 μm x 407 μm window, which was attributed to a diminishing region of activation (Michelson et al., 2019). Their study showed that changes began at regions distal from the stimulated site when delivering stimuli with frequencies greater than 10 Hz. These results are consistent with our study across the larger spatial field (4×4 mm) of the Utah array, as evoked spikes were more likely to be diminished with higher frequency of stimulation at distances closer to the stimulated site. A large difference we observed was the timecourse of the changes; Michelson et al. reported the changes occurred within seconds and plateaued whereas we observed changes occurring for up to 20 minutes. Additionally, we also observed that in some units the probability of evoking a spike increased over time even at longer distances and higher frequencies, which cannot be fully explained by a diminishing region of activation.

Although we did not explicitly measure the duration of changes induced by repetitive stimulation, we observed that they typically lasted less than 2 minutes. Due to the short-lived nature of the induced changes, various mechanisms of short-term synaptic plasticity such as vesicle depletion and facilitation by calcium influx (Citri & Malenka, 2008; Zucker & Regehr, 2002) may best explain our results. Evidence suggests that different forms of short-term plasticity exist for synaptic connections between different cell-types (Blackman et al., 2013; Losonczy et al., 2002). Beyond these differences, previous *in vitro* work by Markram showed that synaptic connections between pyramidal neurons of the same morphological class and interneurons had similar facilitating and depressing characteristics, but with different time courses (Markram et al., 1998).

Together, these results may explain why the frequency-dependent changes that we measured were different for each unit. Although we did not discern any differences between regular and fast spiking neurons for any measure, there are limitations in such cell type classifications with extracellular recordings. Furthermore, we also observed changes in the latency of evoked spikes due to repetitive stimulation, which has previously been shown to occur in the presence of short-term plasticity (Boudkkazi et al., 2007). The latencies typically changed more often with higher frequency stimulation in spikes closer to the stimulated site, similar to the evoked spike probability and inhibitory response changes. Future studies with specific differentiation between cell and synapse types may shed more light on whether cell-type specific differences account for the variability across spikes.

### Excitation and inhibition are independently activated but modulated together within an interconnected network

The balance between excitation and inhibition within the cortex is a much-studied topic and is highly relevant to neural computation. Though the examined network size, location, and synaptic connections vary greatly, the strong consensus is that excitation and inhibition are generally comodulated (Chen, 2004; Haider, 2006; Isaacson & Scanziani, 2011; Rubin et al., 2017; Turrigiano, 2012; Xue et al., 2014). Whether a stimulus evoked a spike on a trial-by-trial basis did not affect the subsequent inhibitory response, but we found that the probability of evoking a spike and the duration of inhibition were frequently positively or negatively correlated over time. Units close to the stimulated site typically had positively correlated evoked spike probability and inhibition duration, both of which were negatively correlated with firing rate. Units far from the stimulation channel, however, had positively correlated evoked spike probabilities and firing rates, which were both negatively correlated with inhibition duration.

The effect of distance can be explained by the fact that sites closer to stimulation are more likely to be activated by ICMS (Butovas & Schwarz, 2003; Hao et al., 2016). Due to the feedforward and feedback inhibitory circuitry, if the total excitation increased or decreased due to short-term plasticity, the inhibition should change in a positively correlated manner. Sites far from stimulation, however, are not activated as comprehensively and are thus less likely to be susceptible to short-term plasticity. Similarly, the negative correlations in evoked spike probability and inhibition at these far sites are likely due to network dynamics, whereas the positive correlations in closer sites are likely due changes in short-term plasticity caused by direct activation via ICMS.

## Acknowledgements

We thank Larry Shupe for programming and software support and Rebekah Schaefer for assistance with animal care, handling, training, and surgery. We also thank Irene Rembado and Nikolai Dembrow for helpful discussion. This work was supported by the National Institutes of Health (NS012542, RR00166, and NS118781) and the National Science Foundation (EEC-1028725).

## Notes

### Competing Interest Statement

The authors have declared no competing interest.

### Summary of Updates

All figures revised to show distributions, included covarying evoked spikes.

## References

Abbott, L. F., & Regehr, W. G. (2004). Synaptic computation. Nature, 431(October), 796–803.

Adesnik, H., & Scanziani, M. (2010). Lateral competition for cortical space by layer-specific horizontal circuits. Nature, 464(7292), 1155–1160.

Berman, N. J., Douglas, R. J., Martin, K. A. C., & Whitteridge, D. (1991). Mechanisms of inhibition in cat visual cortex. Journal of Physiology, 440, 697–722.

Bettler, B., Kaupmann, K., Mosbacher, J., & Gassmann, M. (2004). Molecular structure and physiological functions of GABAB receptors. Physiological Reviews, 84(3), 835–867.

Blackman, A. V., Abrahamsson, T., Costa, R. P., Lalanne, T., & Sjöström, P. J. (2013). Target-cell-specific short-term plasticity in local circuits. Frontiers in Synaptic Neuroscience, 5(DEC), 1–13.

Borchers, S., Himmelbach, M., Logothetis, N., & Karnath, H. (2012). Direct electrical stimulation of human cortex — the gold standard for mapping brain functions? Nature Reviews Neuroscience, 13(January), 63–70.

Boudkkazi, S., Carlier, E., Ankri, N., Caillard, O., Giraud, P., Fronzaroli-Molinieres, L., & Debanne, D. (2007). Release-dependent bariations in synaptic latency: A putative code for short- and long-term synaptic dynamics. Neuron, 56(6), 1048–1060.

Butovas, S., Hormuzdi, S. G., Monyer, H., & Schwarz, C. (2006). Effects of electrically coupled inhibitory networks on local neuronal responses to intracortical microstimulation. Journal of Neurophysiology, 96(3), 1227–1236.

Butovas, S., & Schwarz, C. (2003). Spatiotemporal rffects of microstimulation in rat neocortex: A parametric study using multielectrode recordings. Journal of Neurophysiology, 90(5), 3024–3039.

Chen, R. (2004). Interactions between inhibitory and excitatory circuits in the human motor cortex. Experimental Brain Research, 154(1), 1–10.

Citri, A., & Malenka, R. C. (2008). Synaptic plasticity: Multiple forms, functions, and mechanisms. Neuropsychopharmacology, 33(1), 18–41.

Connors, B. W., & Gutnick, M. J. (1990). Intrinsic firing patterns of diverse neocortical neurons. Trends in Neurosciences, 13(3), 99–104.

Connors, B. W., Malenka, R. C., & Silva, L. R. (1988). Two inhibitory postsynaptic potentials, and GABAA and GABAB receptor-mediated responses in neocortex of rat and cat. The Journal of Physiology, 406(1), 443–468.

Cruikshank, S. J., Lewis, T. J., & Connors, B. W. (2007). Synaptic basis for intense thalamocortical activation of feedforward inhibitory cells in neocortex. Nature Neuroscience, 10(4), 462–468.

Dadarlat, M. C., Sun, Y., & Stryker, M. P. (2019). Widespread activation of awake mouse cortex by electrical stimulation. International IEEE/EMBS Conference on Neural Engineering, NER, 2019-March, 1113–1117.

English, D. F., McKenzie, S., Evans, T., Kim, K., Yoon, E., & Buzsáki, G. (2017). Pyramidal cell-interneuron circuit architecture and dynamics in hippocampal networks. Neuron, 96(2), 505–520.e7.

Flesher, S. N., Collinger, J. L., Foldes, S. T., Weiss, J. M., Downey, J. E., Tyler-Kabara, E. C., Bensmaia, S. J., Schwartz, A. B., Boninger, M. L., & Gaunt, R. A. (2016). Intracortical microstimulation of human somatosensory cortex. Science Translational Medicine, 8(361), 1–11.

Griffin, D. M., Hudson, H. M., Belhaj-Saif, A., & Cheney, P. D. (2011). Hijacking cortical motor output with repetitive microstimulation. Journal of Neuroscience, 31(37), 13088–13096.

Gustafsson, B., & Jankowska, E. (1976). Direct and indirect activation of nerve cells by electrical pulses applied extracellularly. The Journal of Physiology, 258(1), 33–61.

Haider, B. (2006). Neocortical network activity in vivo is generated through a dynamic balance of excitation and inhibition. Journal of Neuroscience, 26(17), 4535–4545.

Hao, Y., Riehle, A., & Brochier, T. G. (2016). Mapping horizontal spread of activity in monkey motor cortex using single pulse microstimulation. Frontiers in Neural Circuits, 10(December), 1–16.

Hartmann, K., Thomson, E. E., Zea, I., Yun, R., Mullen, P., Canarick, J., Huh, A., & Nicolelis, M. A. L. (2016). Embedding a panoramic representation of infrared light in the adult rat somatosensory cortex through a sensory neuroprosthesis. Journal of Neuroscience, 36(8).

Helmstaedter, M., Staiger, J. F., Sakmann, B., & Feldmeyer, D. (2008). Efficient recruitment of layer 2/3 interneurons by layer 4 input in single columns of rat somatosensory cortex. Journal of Neuroscience, 28(33), 8273–8284.

Histed, M. H., Bonin, V., & Reid, R. C. (2009). Direct activation of sparse, distributed populations of cortical neurons by electrical microstimulation. Neuron, 63(4), 508–522.

Hussin, A. T., Boychuk, J. A., Brown, A. R., Pittman, Q. J., & Campbell Teskey, G. (2015). Intracortical microstimulation (ICMS) activates motor cortex layer 5 pyramidal neurons mainly transsynaptically. Brain Stimulation, 8(4), 742–750.

Isaacson, J. S., & Scanziani, M. (2011). How inhibition shapes cortical activity. Neuron, 72(2), 231–243.

Jackson, A., & Fetz, E. E. (2011). Interfacing with the computational brain. IEEE Transactions on Neural Systems and Rehabilitation Engineering.

Kara, P., Pezaris, J. S., Yurgenson, S., & Reid, R. C. (2002). The spatial receptive field of thalamic inputs to single cortical simple cells revealed by the interaction of visual and electrical stimulation. 99(25), 1–6.

Klink, P. C., Dagnino, B., Gariel-Mathis, M. A., & Roelfsema, P. R. (2017). Distinct feedforward and feedback effects of microstimulation in visual cortex reveal neural mechanisms of texture segregation. Neuron, 95(1), 209–220.e3.

Kumar, S. S., Wülfing, J., Okujeni, S., Boedecker, J., Riedmiller, M., & Egert, U. (2016). Autonomous optimization of targeted stimulation of neuronal networks. PLoS Computational Biology, 12(8), 1–22.

Lebedev, M. A., & Nicolelis, M. A. L. (2017). Brain-machine interfaces: From basic science to neuroprostheses and neurorehabilitation. Physiological Reviews, 97(2), 767–837.

Lesser, R. P., Lee, H. W., Webber, W. R. S., Prince, B., Crone, N. E., & Miglioretti, D. L. (2008). Short-term variations in response distribution to cortical stimulation. Brain, 131(6), 1528–1539.

Logothetis, N. K., Augath, M., Murayama, Y., Rauch, A., Sultan, F., Goense, J., Oeltermann, A., & Merkle, H. (2010). The effects of electrical microstimulation on cortical signal propagation. Nature Neuroscience, 13(10), 1283–1291.

Losonczy, A., Zhang, L., Shigemoto, R., Somogyi, P., & Nusser, Z. (2002). Cell type dependence and variability in the short-term plasticity of EPSCs in identified mouse hippocampal interneurones. Journal of Physiology, 542(1), 193–210.

Markram, H., Wang, Y., & Tsodyks, M. (1998). Differential signaling via the same axon of neocortical pyramidal neurons. Proceedings of the National Academy of Sciences of the United States of America, 95(9), 5323–5328.

Matsumura, M., Chen, D., Sawaguchi, T., Kubota, K., & Fetz, E. E. (1996). Synaptic interactions between primate precentral cortex neurons revealed by spike-triggered averaging of intracellular membrane potentials in vivo. Journal of Neuroscience, 16(23), 1–11.

McCormick, D. A., Connors, B. W., Lighthall, J. W., & Prince, D. A. (1985). Comparative electrophysiology of pyramidal and sparsely spiny stellate neurons of the neocortex. Journal of Neurophysiology, 54(4), 782–806.

McIntyre, C. C., & Grill, W. M. (2000). Selective microstimulation of central nervous system neurons. Annals of Biomedical Engineering, 28(3), 219–233.

Michelson, N. J., Eles, J. R., Vazquez, A. L., Ludwig, K. A., & Kozai, T. D. Y. (2019). Calcium activation of cortical neurons by continuous electrical stimulation: Frequency dependence, temporal fidelity, and activation density. Journal of Neuroscience Research, 97(5), 620–638.

Ranck, J. B. (1975). Which elements are excited in electrical stimulation of mammalian central nervous system: A review. Brain Research, 98(3), 417–440.

Rubin, R., Abbott, L. F., & Sompolinsky, H. (2017). Balanced excitation and inhibition are required for high-capacity, noise-robust neuronal selectivity. Proceedings of the National Academy of Sciences of the United States of America, 114(44), E9366–E9375.

Rudy, B., Fishell, G., Lee, S., & Hjerling-Leffler, J. (2013). Three groups of interneurons account for nearly 100 % of Neocortical GABAergic Neurons. Dev Neurobiol, 71(1), 45–61.

Sejnowski, T. J., Churchland, P. S., & Movshon, J. A. (2014). Putting big data to good use in neuroscience. Nature Neuroscience, 17(11), 1440–1441.

Shupe, L. E., Miles, F. P., Jones, G., Yun, R., Mishler, J., Rembado, I., Murphy, R. L., Perlmutter, S. I., & Fetz, E. E. (2021). Neurochip3: An autonomous multichannel bidirectional brain-computer interface for closed-loop activity-dependent stimulation. Frontiers in Neuroscience, 15(August), 1–15.

Silberberg, G., & Markram, H. (2007). Disynaptic inhibition between neocortical pyramidal cells mediated by martinotti cells. Neuron, 53(5), 735–746.

Stoney, S. D., Thompson, W. D., & Asanuma, H. (1968). Excitation of pyramidal tract cells by intracortical microstimulation: effective extent of stimulating current. Journal of Neurophysiology, 31(5), 659–669.

Tehovnik, E. J., Tolias, A. S., Sultan, F., Slocum, W. M., & Logothetis, N. K. (2006). Direct and indirect activation of cortical neurons by electrical microstimulation. Journal of Neurophysiology, 96(2), 512–521.

Tremblay, R., Lee, S., & Rudy, B. (2016). GABAergic interneurons in the neocortex: From cellular properties to circuits. Neuron, 91(2), 260–292.

Turrigiano, G. (2012). Homeostatic synaptic plasticity: Local and global mechanisms for stabilizing neuronal function. Cold Spring Harbor Perspectives in Biology, 4(1), a005736.

Weihberger, O., Okujeni, S., Mikkonen, J. E., & Egert, U. (2013). Quantitative examination of stimulus-response relations in cortical networks in vitro. Journal of Neurophysiology, 109(7), 1764–1774.

Xue, M., Atallah, B. V., & Scanziani, M. (2014). Equalizing excitation-inhibition ratios across visual cortical neurons. Nature, 511(7511), 596–600.

Zhu, J., Jiang, M., Yang, M., Hou, H., & Shu, Y. (2011). Membrane potential-dependent modulation of recurrent inhibition in rat neocortex. PLoS Biology, 9(3).

Zucker, R. S., & Regehr, W. G. (2002). Short-term synaptic plasticity. Annual Review of Physiology, 64(4), 355–405.

